# A workbench for the translational control of gene expression

**DOI:** 10.1101/2020.01.28.923219

**Authors:** Angelo Valleriani, Davide Chiarugi

## Abstract

Ribosome profiling (Ribo-seq profiling) is the most advanced tool to study the translational control of gene expression. Unfortunately, the resolution of this cutting edge technique is severely limited by a low signal to noise ratio. To tackle this issue, we introduce here a newly designed statistical method for the identification of reproducible Ribo-seq profiles. In the case of *E. coli*, the analysis of 2238 Ribo-seq profiles across 9 independent datasets revealed that only 11 profiles are significantly reproducible. A subsequent data quality check led us to identify one outgroup dataset. By ruling it out, the number of highly reproducible profiles could be raised to 49. Despite its surprisingly small size, this set represents a reliable workbench to both assess the quality of the data and study the factors that influence the translation process.

## I. Introduction

Ribosome profiling (Ribo-seq profiling) is designed to investigate mRNA translation with single nucleotide resolution. It consists in the quick blockage of the translation process in living cells followed by the nuclease mediated digestion of the mRNA not covered by ribosomes [1, 2]. The remaining mRNA fragments, called Ribosome Protected Fragments (RPFs), are deep sequenced and the resulting oligonucleotides (Ribo-seq *reads*) are mapped on the reference mRNAs to obtain a snapshot that captures the position of the ribosomes on the mRNA templates when the translation was blocked. Local differences in the density of the RPFs along the mRNAs reflect the different amount of time spent by the ribosomes in translating each part of the template molecule: the slower the ribosomes on a given position, the more the RPFs will accumulate on that area. Therefore, each Open Reading Frame (ORF) can be associated to a specific *Ribo-seq profile* (**Fig. 1**), *i.e*., a histogram that counts the number of reads that cover each nucleotide. Possible “peaks” and “valleys” in the Ribo-seq profile correspond to slow and fast translated regions, respectively. Events that can affect the local translation dynamics (such as, *e.g*., tRNA modifications, changes in the relative tRNAs concentrations, modifications of ribosomes, and codon mutations or reallocations) would be detected by Ribo-seq and will leave a characteristic signature in the Ribo-seq profiles.

**Figure 1:**
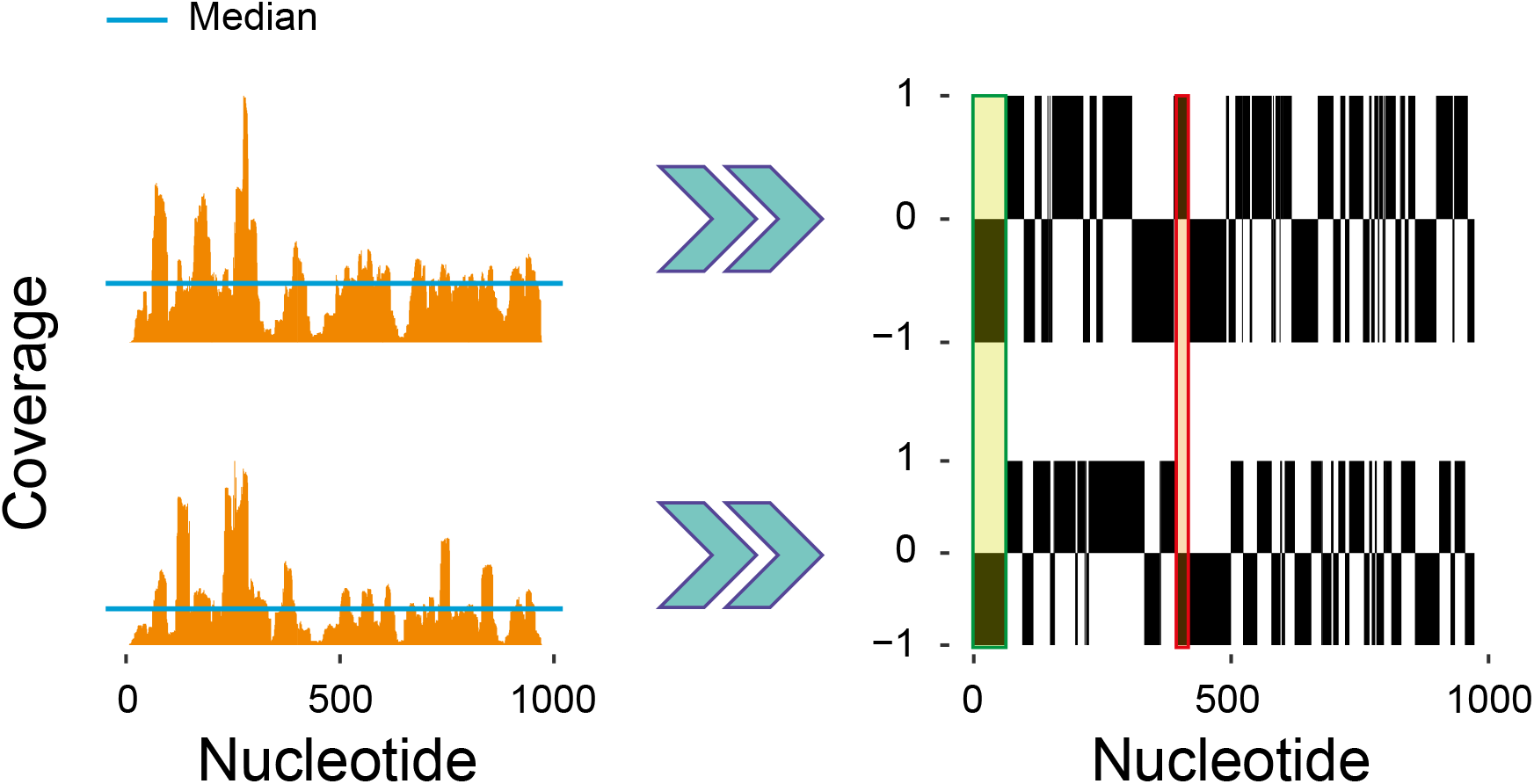
Pairwise comparison of two Ribo-seq profiles of the ispB gene. Two independent Ribo-seq profiles (left), obtained by computing the coverage at each nucleotide position within the ORF are compared to the median coverage to produce the digital ±1 profiles on the right. The digital profiles can be easily compared to detect matches (e.g. green rectangle) and mismatches (e.g. red rectangle). The ratio between the number of matches and the total number of nucleotides in the ORF gives the matching score. A score equal to one means perfect match between the two profiles, whereas a score equal to one-half means poor matching.

Ribo-seq profiling has the unique potential of providing fundamental insight into the translational control of gene expression [2, 3, 1]. Unfortunately, the exploitation of the full power of the technique is severely hampered by the low signal to noise ratio. This issue has a direct impact on the reproducibility of Ribo-seq. Indeed, while the total number of reads per mRNA correlates well across independent experiments [4, 5], a growing body of evidence is suggesting that the experimental reproducibility of the Ribo-seq profiles is, in general, poor [4, 6, 7, 8]. Indeed, multiple variables can affect the data mainly due to the inherent complexity of both the experimental protocol and the data analysis process. For instance, it has been clearly shown that the use and concentrations of translation inhibitors [9], the choice of either the nuclease for hydrolysing the RNA non protected by the ribosomes [10] or the methods for monosome purification or rRNA depletion [11], can lead to different outcomes. From a data analysis perspective, instead, different approaches have provided contradictory conclusions. In this perspective it is worth recalling the case of two independent studies [12, 13] in which the analysis of the same Ribo-seq dataset lead to opposite conclusions about the dependence of the translation rate from Shine-Dalgarno like sequences detected within the considered ORFs.

Here we present a novel data analysis method that allows us to address the aforementioned limitations of Ribo-seq and to recover, when possible, its full resolution. Different from previous work [14, 15], our strategy does not ground on signal processing techniques. Rather, we rely on a data-driven approach to identify and select those reproducible Ribo-seq profiles that emerge from the comparison of independent Ribo-seq experiments performed in different laboratories under the same conditions. The selected profiles will be characterised by an extremely high resolution because the underlying signal will be so sharp that it will overcome the biological noise. These significantly reproducible profiles build a library that can be used as a reference for comparative experiments aimed at detecting differential translation events.

## II. Methods

The fundamental strategy underlying our approach consists in the identification of high resolution Ribo-seq profiles through the systematic comparison of Ribo-seq datasets referring to experiments performed independently in different laboratories and in different time periods. The ORF-specific Ribo-seq profiles that will exhibit a significant degree of similarity in spite of the experimental variability will also bear the sharpest signal and will be, then, selected.

We start by illustrating our workflow with the comparison of two profiles from two independent datasets. In the first step of our workflow, the Ribo-seq profiles (**Fig. 1**, left side) are digitalised by comparing the profile heights at each nucleotide position (*coverage*) with its median value computed along the entire ORF. The result is one *digital Ribo-seq profile* (**Fig. 1**, right side) for each Ribo-seq profile. The pairwise comparison of two digital profiles results in areas of match (**Fig. 1**, right side, green rectangle) and areas of mismatch (**Fig. 1**, right side, red rectangle). Calculating the relative number of matches (the ratio between the number of matches and the length of the ORF in nucleotides) yields the *matching score*.

Intuitively, a matching score close to one could indicate a high degree of similarity between a pair of digitalised profiles, whereas a score around one-half could mean a very poor overlap because the observed matches are likely to have occurred by chance. The extent to which a given matching score denotes possible profiles similarity will be decided through a statistical test that evaluates the score in the context of a true null hypothesis. Here, the null hypothesis is that the score is obtained by chance and, thus, the detected profiles similarities originate from random fluctuations. This concept finds its mathematical counterpart into a null distribution that describes the probability for a matching score to be obtained by chance. The null distribution is created through a data-driven approach: for any Ribo-seq profile involved in a pairwise comparison, a large number of randomised profiles is generated by re-distributing the reads in random positions on the reference ORF. The randomised profiles are, in turn, compared pairwise yielding a large number of random matching scores that build each null distribution. Mapping the matching score on the null distribution will yield one *p*-value (**Figs. 2** and **3**). If the *p*-value will result below a given threshold, the compared Ribo-seq profiles will exhibit a significant degree of similarity. To highlight which specific regions within the Ribo-seq profiles are similar to each other a *consensus sequence* (**Fig. 4**) can be built. The consensus sequence is a character string representing the nucleotides of the reference ORF and coloured in red in those positions where a peak is present in both the profiles (*i.e*. the digitalised profiles values are +1 and the ribosome proceeds slower) and in green where a valley is located. The black color, instead, will be used in all other cases.

**Figure 2:**
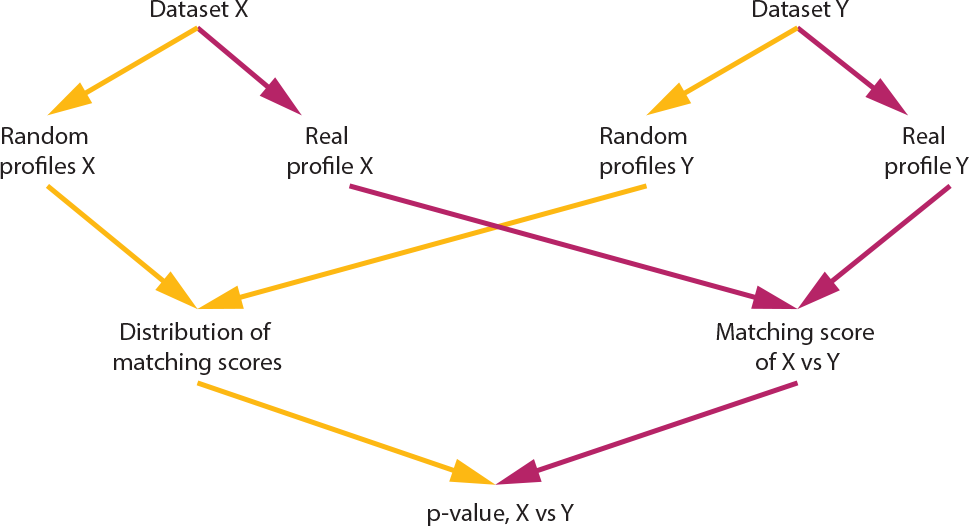
Workflow for the derivation of the matching score significance. For each gene, two Ribo-seq profiles from two independent studies are first digitalised and compared to derive the matching score (red arrows). The RPFs of each dataset are then used to generate random profiles, which are then digitalised and compared pairwise to obtain a distribution of random matching scores (yellow arrows). The real matching score is then mapped on the distribution of random matching scores to derive a p-value (**Fig. 3**). This procedure is repeated for each pair of independent studies and for each gene.

**Figure 3:**
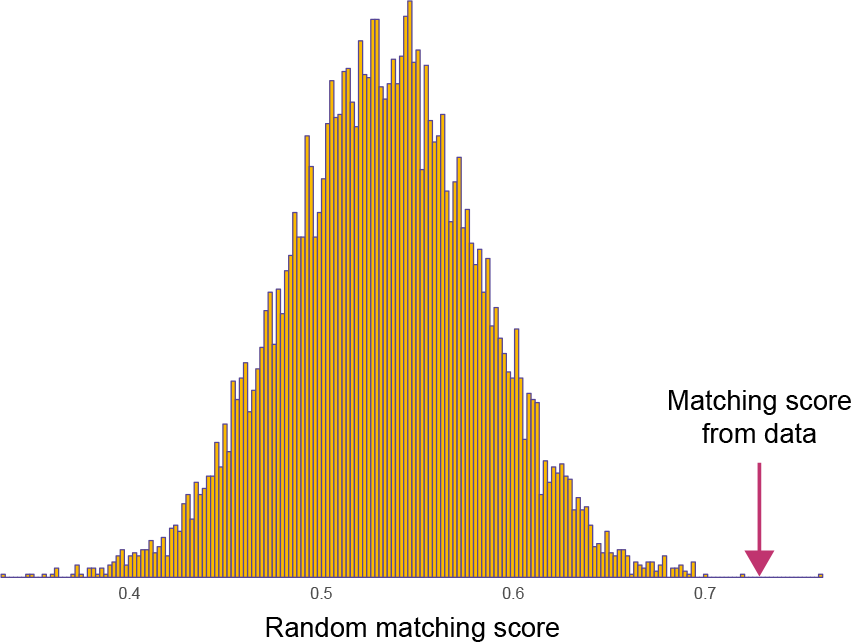
Determination of the p-value from the null distribution. Based on the procedure described in our workflow (**Fig. 2**) the distribution of random scores is generated for each gene and each pair of independent datasets. The score from data (the real score) is compared with the distribution and a p-value is computed as the probability that a score equal or larger than the real score could emerge from the randomisation procedure. A detailed description of the procedure is provided in the Supplementary Section S2.1.

**Figure 4:**
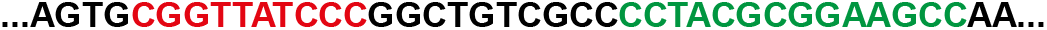
A consensus sequence of fast and slow translation regions. Part of a consensus sequence (computed for rsxC) indicating the nucleotides situated within fast (green) and slow (red) translation regions. The entire set of consensus sequences is presented in the accompanying Supplementary File AllSequences49.xlsx

The core workflow can be scaled up in the typical case when many datasets are considered. The details of the workflow regarding this scenario are reported in the Supplementary Materials. In general, each dataset will contain the information needed to compute one Ribo-seq profile for each protein coding gene of the studied organism. Considering *n* datasets will result in *N* = *n*(*n* − 1)/2 pairwise comparisons for each ORF and, thus, in *N* matching scores. If each dataset will contain the information for an amount *O* of ORFs, a total of *M_TOT_* = *N* × *O* matching scores will be obtained and a set of *M_TOT_* null distributions (one for each pairwise comparison) will be generated. The mapping of the *M_TOT_* matching scores on the correspondent null distributions will yield *N p*-values per ORF. If all the *N* ORF-specific *p*-values will result below a given threshold, the Ribo-seq profiles for that ORF will exhibit a high degree of reproducibility across all datasets and will be selected as part of the high resolution library. The correspondent consensus sequence (**Fig. 4**) will be, then, generated. In this case, the green colour will mark those positions in which at least 75% of the Ribo-seq profiles have a valley whereas the elements coloured in red will indicate a peak in 75% of the profiles. The threshold for the *p*-values is computed for each ORF relying on the Benjamini-Hochberg (BH) procedure that outputs a value which is corrected for possible biases arising from multiple comparisons. The overall sensitivity of our method can be tuned at this stage by choosing more or less conservative False Discovery Rate (FDR) boundaries.

## III. Results

We performed a large-scale analysis considering the *E. coli* Ribo-seq data stored in the GEO repository [16, 17]. Currently, this database hosts 14 collections of dataset (series) that include at least one group of data (sample) from Ribo-seq experiments performed on *E. coli* in various conditions according to the most used experimental protocol [4]. Table S1 summarizes the main features of these series and of the samples contained therein. We clustered this data into homogeneous groups taking into account three main experimental variables, namely the strain’s genotype, the culture medium and the experimental conditions (Table S1). Our analysis regarded a subset of 32 samples that refer to experiments performed culturing wild type *E. coli* strains under control conditions. We partitioned these samples according to the used growth medium and the genotype of the wild types. In this way, we obtained the 7 homogeneous groups (labelled from A to G) reported in Table S2. Each group is composed of samples characterized by combinations of *E. coli* genotype and growth medium and taken from different datasets (GEO Series), *i.e*., performed in different laboratories and time periods. We started the analysis with the largest group A, which is composed of 17 samples collected from 9 different series and obtained from *E. coli* K-12 MG1655 cultured in a MOPS-based medium. Subsequently, using group A as a benchmark, we compared it to the remaining 6 groups (C to G). In this way we evaluated the impact of either the culture medium or the choice of the wild-type’s genotype.

Several samples contain two or more technical replicates, which are known to be more similar to each other than data produced independently from different labs. Thus, initially we considered one sample for each series, putting aside the data referring to replicates that will be later on cast into play. Putting it all together, we consider 9 samples, each belonging to different series. The comparison follows the steps outlined in the Methods and described in detailed in the Supplementary Materials.

Following this strategy, we found that, out of 2238 genes that are in common to all 9 series, the 11 genes listed in Table 1 have significantly reproducible Ribo-seq profiles. For one of these genes, namely rsxC (EG13935), we show the profiles as an illustrative example (**Fig. 5**).

**Figure 5:**
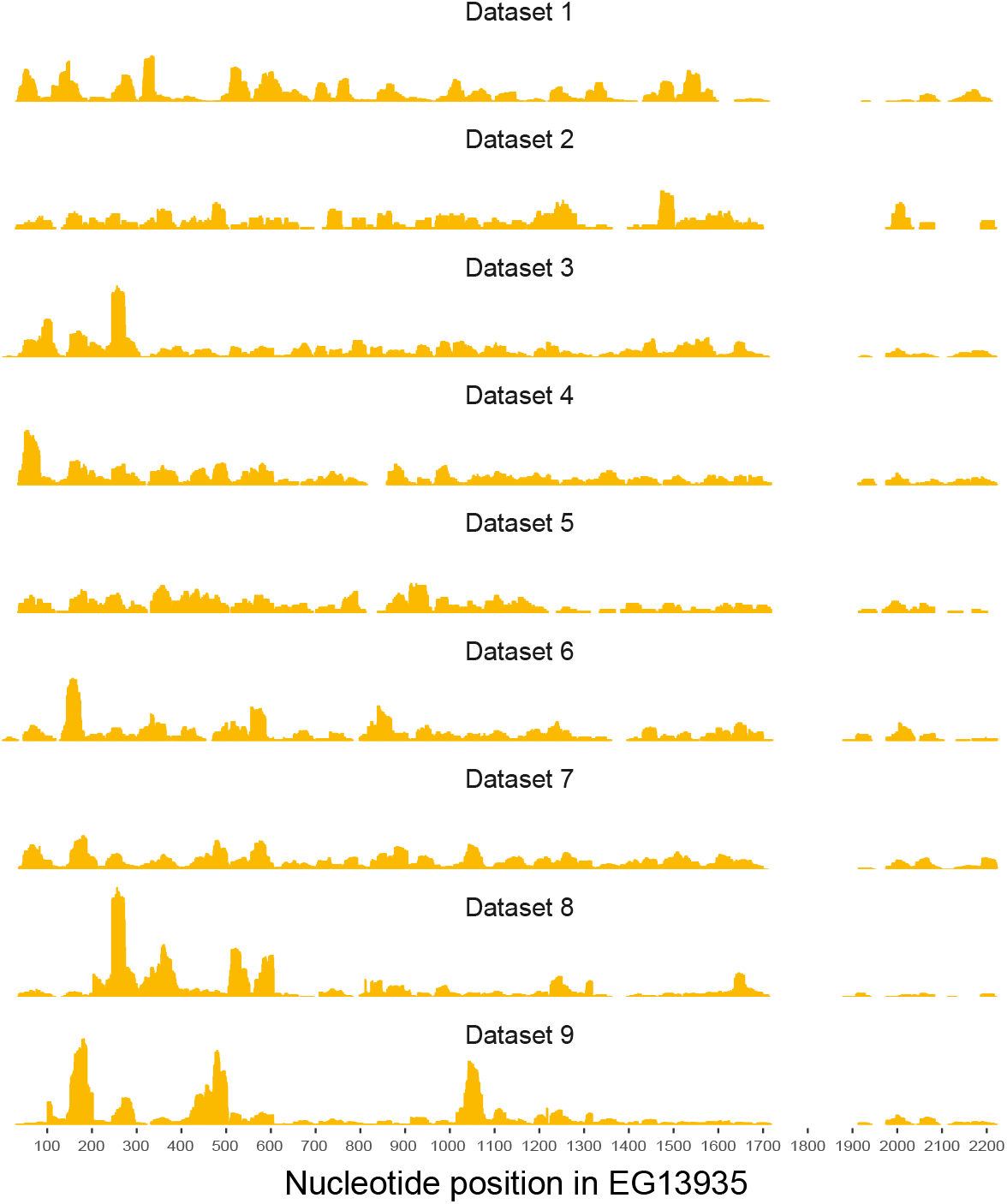
Illustrative example of a significantly reproducible Ribo-seq profile (gene rsxC, EG13935). Note that the comparison takes into account both the areas where the coverage is high and the areas where the coverage is low. Performing by eye a comparative analysis of these profiles would be readily impossible. The entire collection of reproducible profiles is presented in the accompanying Supplementary File AllProfiles49.zip.

**Table 1:**
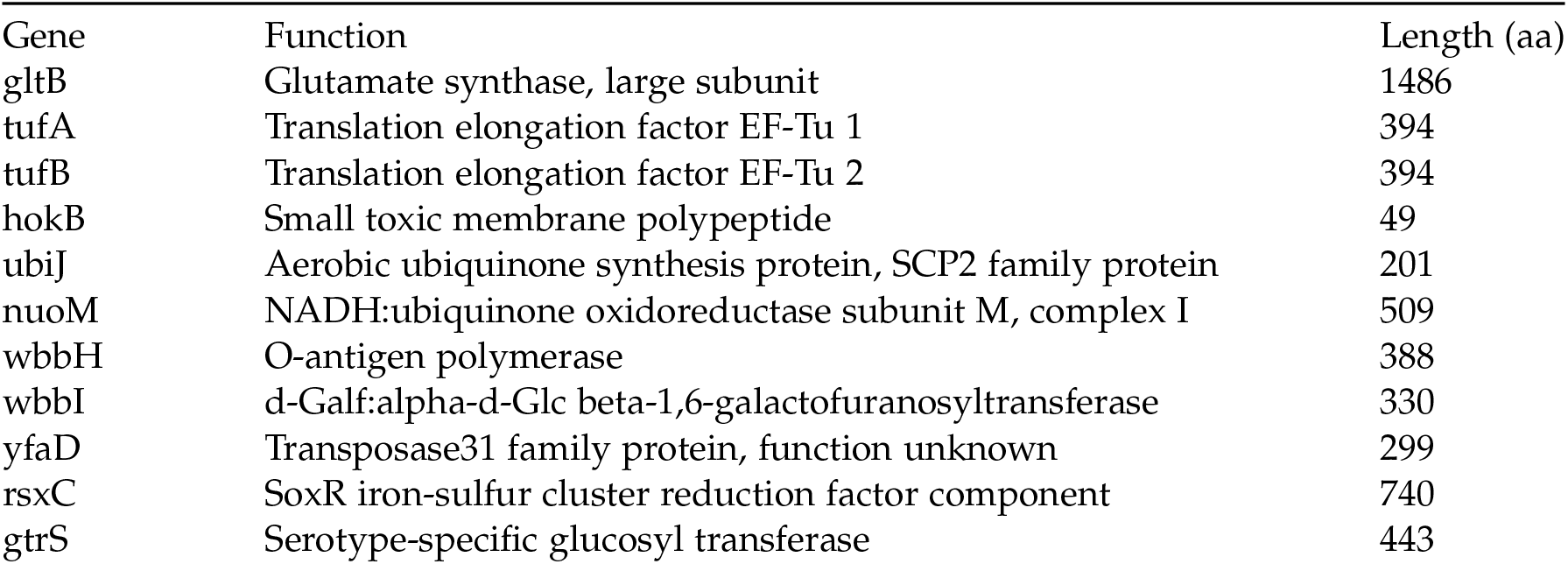
Genes with significantly reproducible Ribo-seq profiles

To check whether the analysis of the replicates of the considered samples might yield different results, we repeated the analysis for each of the replicates. It turned out that the choice of the experimental replicates has no influence on this list.

### i. Samples quality check

One by-product of our analysis allows us to check for the presence of samples that, in spite of being collected in the experimental condition characterizing the group, might be affected by unpredictable biases. To do this, we relied on a tailored jackknife approach. Indeed, we repeated 9 times the last step of the reproducibility analysis excluding each time one sample. After this analysis, it turned out that in all the cases in which the Sample A10*ζ* (Series GSE58637[18]) was excluded, the number of reproducible Ribo-seq profiles raised from 11 to 49 whereas the exclusion of all other samples had no effect. We interpreted this result as originating from a peculiarity of this experiments that we could not better clarify. At this point, for each subsequent analysis presented below we have considered the set of 49 genes as benchmark. The list of these genes is presented in Table S5.

### ii. Impact of the culture medium

We used our method to evaluate whether performing Ribo-seq culture media different from those characterising the group A, might affect experimental reproducibility. To do this, we examined all the samples belonging to the groups B–G listed in Table S2 and we compared them with the samples of group A (without considering sample A10*ζ*).

We started this comparative analysis considering group B, which refers to Ribo-seq data collected from *E. coli* K-12 MG1655 (the same genotype as in group A) cultured in the LB medium. Thus, comparing samples from group B with those from group A allows us to evaluate whether the experimental variable culture medium (in this case LB *vs*. MOPS-rich medium) might influence the reproducibility of Ribo-seq. It turned out that all 49 genes obtained from within group A (excluding A10*ζ*) have Ribo-seq profiles that are significantly robust when changing the culture medium from MOPS-rich to LB. This result was also independent of the replicate chosen for the analysis, as shown in Table S6.

Following the same strategy described above, we tested the samples of group C against our benchmark of 49 genes belonging to group A. Here the same *E. coli*’s genotype characterises both groups A and C, which differ for the chosen culture medium (Minimal Medium in Group C instead of MOPS-rich medium in Group A). In this comparative analysis, only the 2214 ORFs in common between the samples in groups A and C were considered. Noteworthy, this set includes all the 49 ORFs from within group A. When the samples C1*α*, C3*β* and the corresponding replicates were challenged against the benchmark, it turned out that only 24 to 26 Ribo-seq profiles exhibited significant reproducibility.

We interpreted this results, summarized in Table S7 observing that the choice of the LB medium instead of the MOPS rich one has no impact on the reproducibility of Ribo-seq. On the other hand, we observe a significant drop in the number of reproducible profiles when changing from MOPS-rich medium to minimal medium, thus indicating that translation under these two growth conditions is significantly differently regulated.

### iii. Impact of genotype

We reiterated our comparative analysis strategy to examine the role of the chosen *E. coli*’s genotype. Indeed, challenging the samples of the groups D, E, F and G with our benchmark of 49 genes slected from group A, we aimed at testing whether choosing the genotypes BW25113, BWDK or MC4100 instead of MG1655 might reduce the number of reproducible Ribo-seq profiles with respect to the 49 ones identified when the benchmark was analysed in isolation. Table S8 summarises the results of this investigation. It turned out that the choice of the genotype is a variable to consider in terms of experimental reproducibility. Indeed, up to our results, while relying on BW25113 or BWDK genotypes do not impact significantly the experimental reproducibility with respect to the MG1655 genotype, when *E. coli* MC4100 (Samples G1*α*, G2*α* and G3*α*) is chosen, the number of reproducible Ribo-seq profiles falls to 28.

### iv. A set of resilient Ribo-seq profiles

Inspecting the results of the comparative analysis described above (summarised in Table S9), where the impact on profiles reproducibility of either the culture medium or the genotype was investigated, a group of 15 Ribo-seq profiles turned out to be reproducible independently on the experimental condition. Studying the features of the genes corresponding to these extremely resilient Ribo-seq profiles would be surely be interesting but we believe it is beyond the scope of this paper. Nevertheless it worths noticing that, due to both their high reproducibility and relative independence on the experimental condition, the 15 profiles we identified represent excellent candidates as references for Ribo-seq profiles normalisation when it is needed for comparison purposes.

## IV. Discussion

Our method allows the characterisation of a library of significantly reproducible, high resolution Ribo-seq profiles. One of the main strength of our approach lies in its statistical core. Indeed, we evaluate the reproducibility relying on a data-driven statistical framework (based on the FDR concept and on the BH method for multiple testing correction) that does not depend on any model defined *a priori*, that might introduce unpredictable biases in the analysis. The only data-independent parameter that characterises our approach is the FDR threshold that we set to 0.01 in the analysis of the *E. coli* case study. In this case, we accepted 1% of the “positive” results (in our case 1% of the reproducible profiles) to be so by chance. We believe this assumption to be a reasonable compromise between a conservative approach and the need of taking into account the biological variability. In any case, we are confident that our results about *E. coli* are robust with respect to the chosen threshold. Indeed, a test that consisted in varying the FDR threshold within an interval ranging from 0.005 and 0.1 revealed that the number of reproducible profiles obtained analysing the Samples of group A was 49, independently from the threshold value.

With respect to the *E. coli* case study itself, we characterised a library of 11 high resolution Ribo-seq profiles. This library can be expanded to 49 entries after the samples quality check. The consequence of these results is twofold. On the one hand, we provide a small but extremely reliable reference set that can be used as a benchmark for comparative studies. On the other hand, our analysis revealed poor reproducibility of Ribo-seq: up to our criteria a maximum of only 49 out of 2238 profiles could be defined reproducible when 8 different datasets were considered. Therefore, while the latter result might call for a thorough revision of the Ribo-seq’s experimental protocol, we resorted with 11 (49) high reproducible profiles which can be studied enjoying the full, single nucleotide resolution of Ribo-seq. In other words, the general viability of each result in terms of translation’s dynamics coming from the analysis of the “peaks” and “valleys” of the high reproducible profiles will be strongly supported by the clear evidence of being independent from the laboratory performing the experimental procedure.

Within the analysis of the *E. coli* case study, we have shown that our analysis approach can be used to support a possible inspection of the Ribo-seq protocols aimed at identifying the experimental variables that are more likely to affect reproducibility. As an example, we found that performing Ribo-seq on different strains of *E. coli* or growing the bacterium in different culture media, might lead to non-comparable results. The same strategy can easily be extended to other experimental variables thus providing a unique, evidence-based framework to aid the experimental design of Ribo-seq.

A possible alternative to detect similarities between two profiles, would be to compare them through a scatter plot and estimate the degree of correlation between the (relative) coverage in each position of the ORF. Strong correlation would denote high level of similarity between the compared profiles at hand and *vice versa*. While this approach might look more straightforward by intuition, it actually exhibits relevant drawbacks. Indeed, it could be difficult to define a threshold for a given correlation coefficient to be a good indicator of similarity. Moreover, depending on the method chosen for computing the regression coefficient, the contribution of each single point in determining the regression might be different, thus affecting our estimation of reproducibility. These drawbacks are not issues for our method that, in addition, allows to deal straightforwardly with multiple comparisons.

In this work, we do not address the possible causes of the homogeneous dynamical behaviour that characterises the 11 (49) profiles of the obtained library: if we see a reproducible peak in a given profile, what is the origin of this peak? Why are the ribosomes slower at that position? This kind of questions could be be addressed by considering multiple, possible concurrent causes such as the interaction between the nascent protein and the ribosome exit tunnel, the strong interaction of particular nucleotide sequences with the ribosome body or the presence of either specific slow codons in the ORF or mRNA secondary structure downstream. We believe that the study of the consensus sequences (**Fig. 4**) underlying the detected peaks and valleys will provide valuable clues. Additionally, we find relevant to stress that according to our method it is possible to decide about the profiles reproducibility only for those cases in which the signal rises consistently above the noise across the considered datasets. Indeed, even though potentially reproducible, those profiles in which the noise is much stronger than the signal will be called non-reproducible in any case, thus leading to rule out possible important pieces of information. Understanding the origin and the causes of fast and slow regions in the reproducible profiles may provide some tools to highlight the presence of significant signals also in those profiles originally considered non-reproducible.

Finally, it is worth mentioning that our method can be straightforwardly applied to each kind of Ribo-seq dataset including those related to organisms such as yeast, mice or humans. In any case the obtained high resolution Ribo-seq profiles libraries will serve as reliable benchmarks to detect possible differential translation events and, thus, to obtain precious insights into the translational control of gene expression.

## References

[1] Ingolia, N. T. (2016) Ribosome Footprint Profiling of Translation throughout the Genome. Cell, 165(1), 22–33.

[2] Brar, G. A. and Weissman, J. S. (2015) Ribosome profiling reveals the what, when, where and how of protein synthesis. Nature Reviews Molecular Cell Biology, 16(11), 651–664.

[3] Jackson, R. and Standart, N. (2015) The awesome power of ribosome profiling. RNA, 21(4), 652–654.

[4] Oh, E., Becker, A. H., Sandikci, A., Huber, D., Chaba, R., Gloge, F., Nichols, R. J., Typas, A., Gross, C. A., Kramer, G., Weissmann, J. S., and Bukau, B. (2011) Selective ribosome profiling reveals the co-translational chaperone action of trigger factor in vivo. Cell, 6(147), 1295–1308.

[5] Sin, C., Chiarugi, D., and Valleriani, A. (2016) Quantitative assessment of ribosome drop-off in E. coli. Nucleic Acids Research, 44(6), 2528–2537.

[6] Calviello, L. and Ohler, U. (2017) Beyond Read-Counts: Ribo-seq Data Analysis to Understand the Functions of the Transcriptome. Trends in genetics: TIG, 33(10), 728–744.

[7] O’Connor, P. B. F., Andreev, D. E., and Baranov, P. V. (2016) Comparative survey of the relative impact of mRNA features on local ribosome profiling read density. Nature Communications, 7, 12915.

[8] Diament, A. and Tuller, T. (2016) Estimation of ribosome profiling performance and reproducibility at various levels of resolution. Biology Direct, 11(1), 24.

[9] Gerashchenko, M. V. and Gladyshev, V. N. (2014) Translation inhibitors cause abnormalities in ribosome profiling experiments. Nucleic Acids Research, 42(17).

[10] Gerashchenko, M. V. and Gladyshev, V. N. (2017) Ribonuclease selection for ribosome profiling. Nucleic Acids Research, 45(2), e6.

[11] Hussmann, J. A., Patchett, S., Johnson, A., Sawyer, S., and Press, W. H. (2015) Understanding Biases in Ribosome Profiling Experiments Reveals Signatures of Translation Dynamics in Yeast. PLoS Genetics, 11(12).

[12] Li, G.-W., Oh, E., and Weissman, J. S. (2012) The anti-Shine-Dalgarno sequence drives translational pausing and codon choice in bacteria. Nature, 484(7395), 538–541.

[13] Mohammad, F., Woolstenhulme, C. J., Green, R., and Buskirk, A. R. (2016) Clarifying the Translational Pausing Landscape in Bacteria by Ribosome Profiling. Cell Reports, 14(4), 686–694.

[14] Calviello, L., Mukherjee, N., Wyler, E., Zauber, H., Hirsekorn, A., Selbach, M., Landthaler, M., Obermayer, B., and Ohler, U. (2016) Detecting actively translated open reading frames in ribosome profiling data. Nature Methods, 13(2), 165–170.

[15] Birkeland, Å., Chyżyńska, K., and Valen, E. (2018) Shoelaces: an interactive tool for ribosome profiling processing and visualization. BMC Genomics, 19(1), 543.

[16] Barrett, T., Wilhite, S. E., Ledoux, P., Evangelista, C., Kim, I. F., Tomashevsky, M., Marshall, K. A., Phillippy, K. H., Sherman, P. M., Holko, M., Yefanov, A., Lee, H., Zhang, N., Robertson, C. L., Serova, N., Davis, S., and Soboleva, A. (nov, 2012) NCBI GEO: archive for functional genomics data sets—update. Nucleic Acids Research, 41(D1), D991–D995.

[17] Edgar, R., Domrachev, M., and Lash, A. E. (jan, 2002) Gene Expression Omnibus: NCBI gene expression and hybridization array data repository.. Nucleic acids research, 30(1), 207–10.

[18] Guo, M. S., Updegrove, T. B., Gogol, E. B., Shabalina, S. A., Gross, C. A., and Storz, G. (2014) MicL, a new σE-dependent sRNA, combats envelope stress by repressing synthesis of Lpp, the major outer membrane lipoprotein. Genes and Development, 28(14), 1620–1634.

