## Supplementary Materials for "A workbench for the translational control of gene expression"

#### S.1 Details about the analysis method

Our analysis of the ORF-specific Ribo-seq profiles consist of two phases, detailed in the following.

##### S.1.1 Upstream phase

The upstream phase allows us to compute the Ribo-seq profiles starting from the raw Ribo-seq data. To reconstruct the Ribo-seq profiles, the reads need to be mapped on the respective reference sequences. To this aim we tailored a bioinformatics software pipeline designed in house for obtaining a high mapping accuracy. For each analysed dataset, our pipeline outputs a specific file reporting the positional information of each read, *i.e.* the 5' and 3' coordinates of each read within the reference genome. This information is used to build the Ribo-seq profiles that represent the input of the subsequent analysis.

##### S.1.2 Downstream phase

The downstream phase is the core of our method as it allows to evaluate the statistical significance of similarity and differences between Ribo-seq profiles coming from different datasets.

###### S.1.2.1 Quantifying similarities between Ribo-seq profiles: a signal digitalisation strategy.

The first step of the downstream phase allows the pairwise comparison of ORF-specific Ribo-seq profiles coming from different datasets. To this aim, our strategy is articulated as follows:

- *Signal digitalisation*: given a Ribo-seq profile, we compute the median of the coverage values at each nucleotide and we assign +1 to the positions having a coverage value higher than the median. We assign $-1$ otherwise. In this way we convert each Ribo-seq profile into the corresponding *digitalised profile* (see Figure S.1 for a graphical representation), *i.e.*, a vector having the length of the associated ORF and containing a sequence of $-1$ and 1. To avoid having values of the profile at *exactly* the same value of the median, before computing the median we add a small noise with mean zero e small variance to each coverage value. In the hypothetical case of a Ribo-seq profile constant along the whole mRNA, adding the small noise would create digital profiles jumping randomly over the values $-1$ and $+1$.

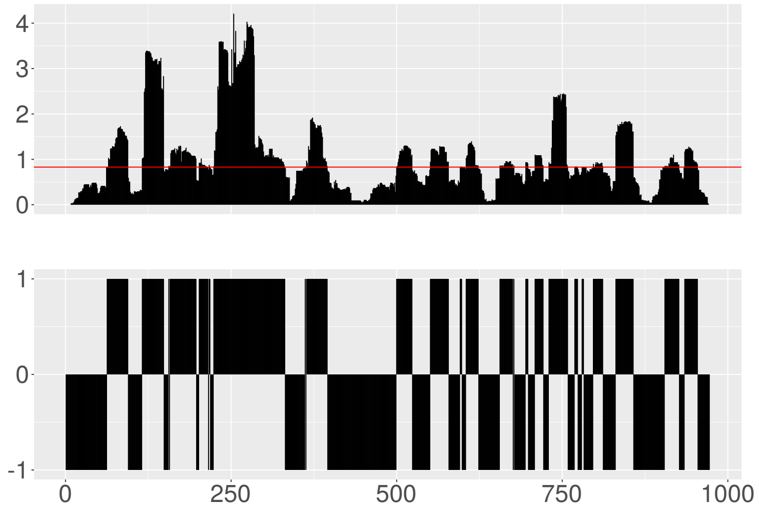


Figure S.1: Example of a Ribo-seq profile (top figure) and the correspondent digitalised profile (bottom figure) for the gene ispB (EG10017) of *E. coli* taken from the Dataset 1 of Table S.2. x-axis: position within the ORF (nucleotides); y-axis (top): relative coverage (number of mapping reads/total number of reads mapping on the ORF.); Red horizontal line (top): median of the Ribo-seq profile; y-axis (bottom): y-coordinate of the digitalised profile ($+1$: the corresponding coverage value is *above* the median; $-1$: the corresponding coverage value is *below* the median.)

- *Comparison of the digital profiles*: we use the digital profiles to quantify the similarities between Ribo-seq profiles coming from different datasets and referring to the same ORF. To this aim, we assigned a *similarity score* to each pairwise comparison. This score was computed aligning each pair of associated digital profiles and counting the number of matches, *i.e.*, the number of times $n$ that the same symbol (either $-1$ or $+1$) appeared in the same position in both profiles, as illustrated in Figure 1 (main text). Then, we divided $n$ by the length (number of nucleotides) of the corresponding ORF, thus obtaining a score ranging from 0.5 (random matches) to 1. The mathematical lowest boundary value for the score is 0. Occasionally, scores with values slightly smaller than 0.5 are observed but in general the comparison between two random and independent profiles would give a score very close to 0.5.

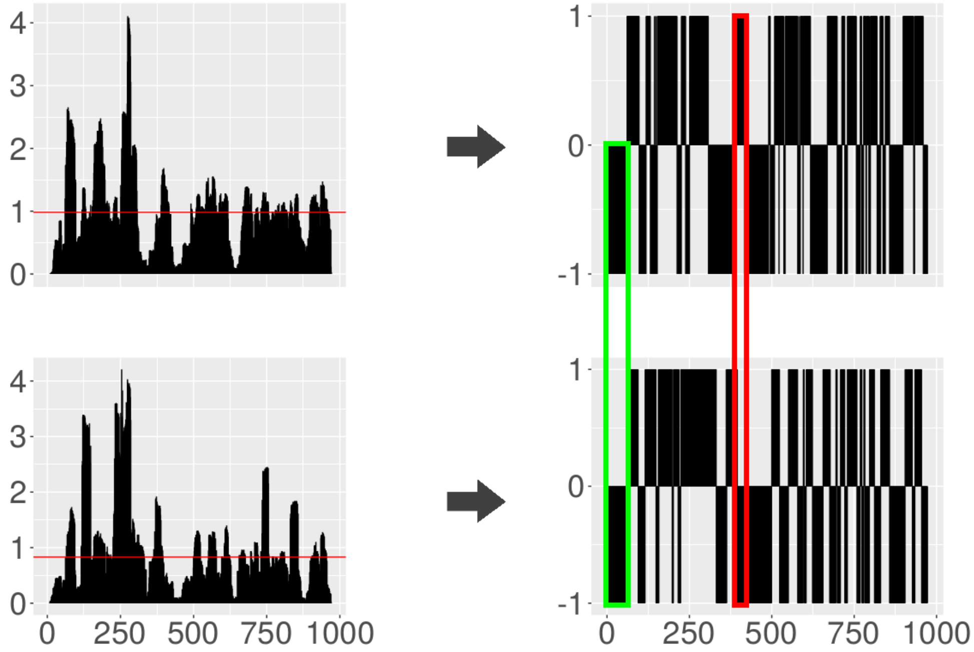


Figure S.2: Pairwise comparison of two Ribo-seq profiles. A pair of digitalised profiles (right side of the figure) is computed from two Ribo-seq profiles (left part of the figure) referring to the same ORF - the ispB gene (EG10017) of *E. coli* - but coming from two different Ribo-seq datasets, namely the Dataset A1$\alpha$ (top-left) and A3$\beta$ (bottom-left) of Table S.2. The green and red rectangles spanning the two digitalised profiles (right side of the Figure) highlight match and mismatch areas respectively. x-axis: position within the ORF (nucleotides); y-axis (left side): relative coverage (number of mapping reads/total number of reads mapping on the ORF.); y-axis (right side): y-coordinate of the digitalised profile ($+1$: the corresponding coverage is *above* the median; $-1$: the corresponding coverage is *below* the median). Red horizontal lines (left side): median coverage.

The similarity score *per se* does not provide any information about the relevance of the amount of matches/mismatches between the compared profiles. In other words, given that each similarity score has a certain probability of being obtained by chance, it is crucial to devise a method to identify those values that indicate statistically significant similarities between two Ribo-seq profiles. To do so, we used the strategy described below.

###### S.1.2.2 A data-driven hypothesis test to assess the reproducibility of Ribo-seq profiles .

Our strategy for assessing the significance of a given similarity score consists of two steps:

*Construction of the null model*: given a pair Ribo-seq profiles referring to the same ORF and coming from two different experiments, *e.g.* the two profiles reported in Figure 1 (main text), the null model ($H_0$) represents the distribution of the similarity scores as they would be if the matches and mismatches between the two profiles are due to randomness. To build such a distribution, we consider the two sets of reads that generated the Ribo-seq profiles in hand and we re-distribute them randomly on the respective ORF, thus generating a pair of *random Ribo-seq profile.* Starting from them and following the procedure used for creating the ORF-specific digitalised profiles it is, then, possible to compute a pair of *digitalised random profiles.* In turn, these profiles can be compared pairwise according to the method explained above thus obtaining a *random similarity score.*

Reiterating this process, we generated $10^4$ pairs of random Ribo-seq profiles and an equal number of digitalised random profiles that, compared pairwise, yielded $10^4$ *random similarity scores.* These scores are, then, used to build a ORF-specific null distribution (Figure S.3) which allows us to estimate the probability of obtaining by chance each similarity score.

It is worth to point out here that our way of building the null distribution through a data-driven random process allows us to formulate the null hypothesis taking solely into account the features of the data without any further hypotheses or approximations.

*Mapping the similarity score on the null distribution*: given a pair of Ribo-seq profiles, the similarity score arising from their comparison is tested for significance by comparing it with the correspondent ORF-specific null distribution, as depicted in Figure S.4. Then, for any given ORF-specific null distribution, a threshold for statistical significance is chosen. If the similarity score is greater than the threshold, then its probability of being obtained by chance is low. We will describe in the following Section our strategy for setting the significance threshold and for

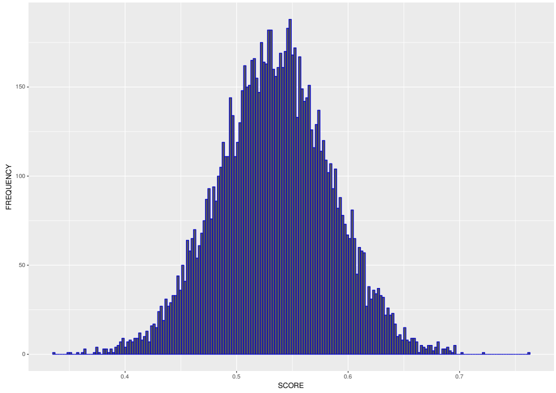


Figure S.3: Null distribution for the gene ispB (EG10017) of *E. coli*. This distribution is obtained comparing pairwise two sets of $10^4$ random digitalised profiles. Each set of random digitalised profiles was computed distributing randomly on the ORF the reads coming from the Samples A1$\alpha$ and A3$\beta$ of Table S.2.

analysing the reproducibility of Ribo-seq profiles in broad scale scenarios, when the analysis involves a large set of ORFs coming from multiple datasets.

#### S.2 Analysing a broad-scale scenario: The *E. coli* case-study.

To illustrate how our method works in a whole organism-scale scenario, we report here the analysis of Ribo-seq profiles referring to *E. coli*. To this aim, we relied on the data stored in the GEO repository [1, 2]. Currently, this database hosts 14 collections of dataset (Series) that include at least one group of data (Sample) resulting from ribo-seq experiments performed on *E. coli* in various conditions according to the most used experimental protocol [3]. Table S.1 summarises the main features of these Series and of the ribo-seq Samples contained therein.

We started our analysis on homogeneous sets of data. To this aim, we clustered the available data into homogeneous groups. This led to 7 homogeneous groups (labelled from A to G) reported in Table S.2. Each group is composed by Samples obtained through experiments characterised by a given combination of *E. coli* genotype and growth medium and taken from different datasets (GEO Series), *i.e.*, performed in different laboratories and time periods. Group A, for instance, is composed of 15 Samples collected from 9 different Series and obtained from *E. coli* k-12 MG1655 cultured in a MOPS-based medium. This group includes the largest amount of data and, thus, we started

| GEO Coordinates (GEO Series ID) | Ribo-seq samples (amount) | Genotype(s) Wild Type(s) | Genotype(s) Mutant(s) | Replicates (amount/sample) | Medium(s) | Stress(es) | Reference |
|---|---|---|---|---|---|---|---|
| GSE64488 | 15 | E. coli k-12 MG1655 | Δefp - ΔepmA/B/C | 2 or None | MOPS, 0.2% glucose or LB | None | [4] |
| GSE90056 | 18 | E. coli k-12 MG1655 | Plasmids added | 3 | MOPS, 0.2% glucose | 42°C×10′; 42°C×20′ | [5] |
| GSE72899 | 5 | E. coli k-12 MG1655 | | 2 or None | MOPS, 1% glucose | None | [6] |
| GSE53767 | 3 | E. coli k-12 MG1655 | | 2 or None | MOPS, 0.2% glucose or MM | None | [7] |
| GSE51052 | 5 | E. coli k-12 MG1655<br>E. coli k-12 BW25113 | Conditional Mutants | None | MOPS, 0.2% glucose | Leu or Ser starvation | [8] |
| GSE58637 | 2 | E. coli k-12 MG165 | Plasmids added | None | MOPS, 0.2% glucose | | [9] |
| GSE77617 | 4 | E. coli k-12 MG165 | Plasmids added<br>Δgcv - dusB-M3 | None | MOPS, 0.2% glucose | None | [10] |
| GSE35641 | 2 | E. coli k-12 MG1655 | Plasmids added<br>E. coli BJW9 | 2 | MOPS, 0.2% glucose | None | [11] |
| GSE61619 | 3 | E. coli k-12 BW25113 | ΔtolC | 2 | LB | Erythromycin or Telithromycin | [12] |
| GSE86536 | 3 | E. coli BWDK | | None | LB | Chloramphenicol or Linezolid | [13] |
| GSE33671 | 4 | E. coli MC4100 | | 3 or None | LB | | [3] |
| GSE85540 | 5 | E. coli k-12 MG1655 | | 3 or 2 | LB | RelE overespression | [14] |
| GSE56372 | 12 | E. coli k-12 MG1655 | Mutant EP61[15] | 2 | MM | Ethanol 40g/l × 10′ or 70′ | [15] |
| GSE88725 | 15 | E. coli k-12 MG1655 | ΔprfB - ΔprfC | 4, 2 or None | MOPS, 0.2% glucose | None | [16] |

Table S.1: Main features of the 14 GEO Series containing at least one Sample resulting from ribo-seq experiments performed according to the most used experimental protocol [3]. Column 1: ID of the GEO Series. Column 2: number of ribo-seq samples contained in the Series; within each Series also non ribo-seq Samples (typically RNA-seq Samples) can be stored. Column 3: features of the *E. coli* genotypes used in the ribo-seq experiments. Column 4: number of replicates of the ribo-seq Samples. Column 5: culture media used in the experiments referring to the considered ribo-seq Samples. Column 6: type of environmental stress possibly introduced in designing some ribo-seq experiments. Column 7: references.

| Group ID | Group's features | | Samples coordinates | | |
|---|---|---|---|---|---|
| | Genotype | Culture's medium | GEO Series ID | GEO Sample ID | Sample ID |
| A | E. coli k-12 MG1655 | MOPS, 0.2% glucose | GSE64488 | GSM1572266 | $1\alpha$ |
| | | | | GSM1572267 | $2\alpha$ |
| | | | GSE90056 | GSM2396722 | $3\beta$ |
| | | | | GSM2396728 | $4\beta$ |
| | | | | GSM2396734 | $5\beta$ |
| | | | GSE72899 | GSM1874188 | $6\gamma$ |
| | | | | GSM1874189 | $7\gamma$ |
| | | | GSE53767 | GSM1300279 | $8\delta$ |
| | | | GSE51052 | GSM1399615 | $9\epsilon$ |
| | | | GSE58637 | GSM1415871 | $10\zeta$ |
| | | | GSE77617 | GSM2055244 | $11\eta$ |
| | | | GSE35641 | GSM872393 | $12\theta$ |
| | | | | GSM872394 | $13\theta$ |
| | | | GSE88725 | GSM2344796 | $14\iota$ |
| | | | | GSM2344797 | $15\iota$ |
| | | | | GSM2344798 | $16\iota$ |
| | | | | GSM2344799 | $17\iota$ |
| B | E. coli k-12 MG1655 | LB | GSE64488 | GSM1572273 | $1\alpha$ |
| | | | | GSM1572275 | $2\alpha$ |
| | | | GSE85540 | GSM2276800 | $3\beta$ |
| | | | | GSM2276801 | $4\beta$ |
| | | | | GSM2276802 | $5\beta$ |
| C | E. coli k-12 MG1655 | Minimal Medium | GSE56372 | GSM1360049 | $1\alpha$ |
| | | | | GSM1360050 | $2\alpha$ |
| | | | GSE53767 | GSM1300280 | $3\beta$ |
| | | | | GSM1300281 | $4\beta$ |
| D | E. coli k-12 BW25113 | MOPS, 0.2% glucose | GSE51052 | GSM1399617 | $1\alpha$ |
| E | E. coli k-12 BW25113 | LB | GSE61619 | GSM1509451 | $1\alpha$ |
| F | E. coli k-12 BWDK | LB | GSE86536 | GSM2305307 | $1\alpha$ |
| G | E. coli k-12 MC4100 | LB | GSE33671 | GSM832607 | $1\alpha$ |
| | | | | GSM832609 | $2\alpha$ |
| | | | | GSM832611 | $3\alpha$ |

Table S.2: The Samples chosen for our analysis among those reported in Table S.1, clustered according to the *E. coli* genotype and culture medium. Column 1: ID labeling the groups. Column 2: genotype and culture media characterising each group. Column 3: samples coordinates, namely the GEO Series ID, the GEO Samples ID and the ID choosen to refer to the various Samples in this work. IDs sharing the same greek letter are replicates of each other.

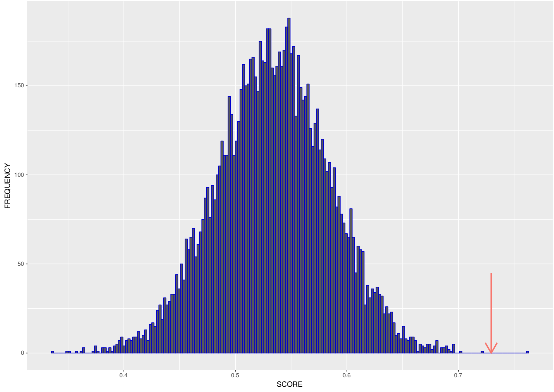


Figure S.4: The null distribution for the gene ispB (EG10017) of *E. coli* depicted in Figure S.3. The red arrow indicates the value of the similarity score obtained comparing the Ribo-seq profiles reported in Figure S.2.

considering it at first. Subsequently, using group A as a benchmark, we compared it to the remaining 6 groups. In this way we have evaluated the impact of either the culture medium or the choice of the wild-type's genotype.

##### S.2.1 Analysis of *E. coli* k-12 MG1655 cultured in MOPS-based medium.

Inspecting column 7 of Table S.2, it can be noticed that some Samples (having the ID which contains the same greek letter) refer to replicates of the same experiment. Within group A, in particular, the couples $1\alpha$ and $2\alpha$, $6\gamma$ and $7\gamma$, $12\theta$ and $13\theta$ as well as the triplet $3\beta$, $4\beta$ and $5\beta$ refer to technical replicates. In our reproducibility analysis, data referring to replicates should be treated differently from the others because it is more likely that the corresponding Samples will be more similar to each other. To this aim, we initially considered one Sample for each Series, putting aside the data referring to replicates that will be later on cast into play. Thus, we started the analysis of group A considering 9 Samples, each belonging to a different Series. More precisely, we compared this data (highlighted in bold in Table S.2) through the following steps:

- *Set up of the coverage matrix*: for this analysis we selected only the 2238 ORFs that have a total coverage greater than 0 in all the 9 considered Samples. For each selected ORF of each selected dataset, we generated a Ribo-seq profile. We stored these profiles in a 2238 $\times$ 9 matrix, named *coverage matrix* (Figure S.5).

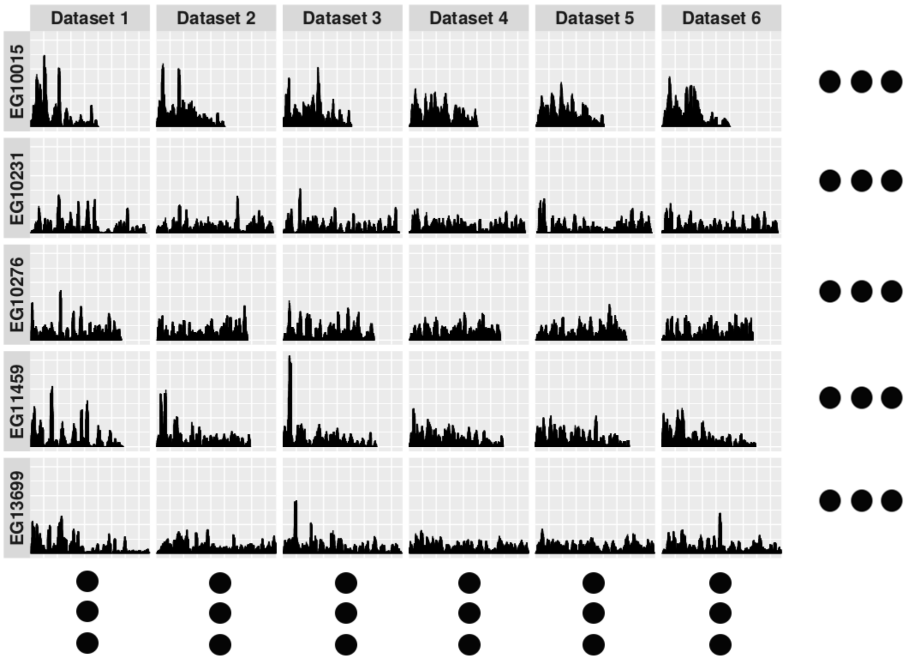


Figure S.5: Representation of the coverage matrix. For the sake of readability only six columns (datasets) and five rows (genes ID) are reported here. The complete matrix is composed by 9 columns and 2238 rows.

- *Elaboration of the digitalised profiles*: following our method, we generated one digitalised profile for each cell of the coverage matrix, obtaining the 2238 $\times$ 9 *digitalised coverage matrix* depicted Figure S.6.

- *Comparison of the digitalised profiles*: For each ORF, i.e. for each row of the digitalised coverage matrix, we compared pairwise all the Ribo-seq profiles coming from the 9 samples at hand. Indeed, we performed 36 - $\binom{9}{2}$ - comparisons for each ORF, obtaining 2238 $\times$ 36 similarity scores, stored in the *scores matrix* (Figure S.7).

- *Assessment of the similarity scores*: the similarity scores provide a quantitative measure of the extent to which a pair of digital profiles (and, thus, the correspondent coverage profiles) match each others. As already noticed, this measure *per se* does not provide enough information about reproducibility, because profile simi-

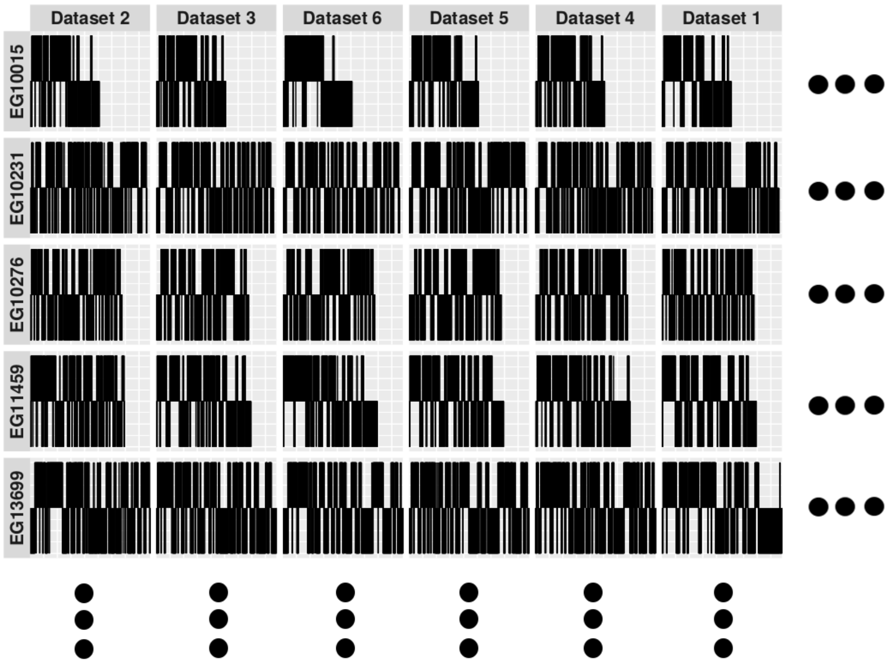


Figure S.6: Representation of the digitalised coverage matrix. For the sake of readability only six columns (datasets) and five rows (genes ID) are reported here. The complete matrix is composed by 9 columns and 2238 rows.

larities might arise by chance. Therefore, we checked the statistical significance of the similarity scores, following the procedure previously described. In particular, for each cell of the score matrix (that is, for each similarity score $s_{i,k}$), we built a distribution of $10^4$ random similarity scores (the null distribution $N_{i,k}$), thus obtaining $2238 \times 36$ null distributions matrix. As expected, each null distribution fits very closely a normal distribution. We take advantage of this fact as follows. For each $s_{i,k}$ contained in the scores matrix and the corresponding null distribution, we computed a z-score $z_{i,k}$, mapping each similarity score on a standard normal distribution through the equation:

$$z_{i,k} = \frac{s_{i,k} - \mu_{N_{i,k}}}{\sigma_{N_{i,k}}} \quad (1)$$

where $mu_{N_{i,k}}$ and $\sigma_{N_{i,k}}$ are, respectively, the mean and standard deviation of the $N_{i,k}$ null distribution. Notice that the mean is not at 0.5. The reason is that when we generate the random profiles, we take into consideration that only RPF's that

| | Dataset 1 vs. Dataset 2 | Dataset 1 vs. Dataset 3 | Dataset 1 vs. Dataset 4 | Dataset 1 vs. Dataset 5 | Dataset 1 vs. Dataset 6 | ··· |
|---|---|---|---|---|---|---|
| **EG10015** | 0.6 | 0.7 | 0.5 | 0.4 | 0.8 | ··· |
| **EG10231** | 0.4 | 0.6 | 0.8 | 0.5 | 0.6 | ··· |
| **EG10276** | 0.9 | 0.8 | 0.5 | 0.6 | 0.9 | ··· |
| **EG11459** | 0.5 | 0.8 | 0.7 | 0.6 | 0.7 | ··· |
| **EG13699** | 0.8 | 0.7 | 0.7 | 0.9 | 0.6 | ··· |
| ··· | ··· | ··· | ··· | ··· | ··· | ··· |
| ··· | ··· | ··· | ··· | ··· | ··· | ··· |
| ··· | ··· | ··· | ··· | ··· | ··· | ··· |

Figure S.7: Representation of the scores matrix. Each column correspond to a pairwise comparison between two datasets. For the sake of readability only 5 columns and 5 rows are reported here. The complete matrix is composed by 36 columns and 2238 rows.

mapped entirely onto the ORF were considered in the analysis of the experimental Ribo-seq data. For this reason, the matching score gets a systematic positive contribution from the regions of the ORF closely downstream of the start codon and closely upstream of the stop codon. This positive contribution shifts the mean to the right of 0.5. We then computed the p-value $p_{i,k}$, as the integral:

$$p_{i,k} = \int_{z_{i,k}}^{+\infty} \mathcal{N}_S(z) dz \tag{2}$$

where $\mathcal{N}_S(z)$ is the standard normal distribution. The results of this process can be summarised into a $2238 \times 36$ matrix (call it *p-values matrix*, Figure S.8) containing all the computed p-values and composed by one column for each pairwise comparison and one row for each considered ORF. Each $p_{i,k}$ quantifies the probability of obtaining a similarity score at least as extreme as the corresponding $s_{i,k}$, given that the null hypothesis is true. In our context, the lower the p-value, the lower the probability that the similarity between the compared pairs of (digital) Ribo-seq profiles occur by chance. In other words, low p-values are more likely to characterise significantly similar (reproducible) Ribo-seq profiles.

- *Identification of the "significantly reproducible" Ribo-seq profiles* Informally, our strategy consists in inspecting each row of the p-values matrix. Then, we define reproducible the Ribo-seq profiles referring to those rows featuring all the p-values below a chosen significance threshold. To cast our strategy into a more rigorous

| | Dataset 1 vs. Dataset 2 | Dataset 1 vs. Dataset 3 | Dataset 1 vs. Dataset 4 | Dataset 1 vs. Dataset 5 | Dataset 1 vs. Dataset 6 | ... |
|---|---|---|---|---|---|---|
| **EG10015** | $8.8 \times 10^{-4}$ | $2.4 \times 10^{-4}$ | $6.1 \times 10^{-4}$ | $2.7 \times 10^{-7}$ | $1.4 \times 10^{-7}$ | ... |
| **EG10231** | $6.4 \times 10^{-3}$ | $2.0 \times 10^{-5}$ | $5.6 \times 10^{-5}$ | $2.5 \times 10^{-4}$ | $8.0 \times 10^{-2}$ | ... |
| **EG10276** | $3.1 \times 10^{-4}$ | $2.9 \times 10^{-3}$ | $8.6 \times 10^{-3}$ | $3.0 \times 10^{-3}$ | $3.8 \times 10^{-1}$ | ... |
| **EG11459** | $4.4 \times 10^{-3}$ | $3.3 \times 10^{-1}$ | $2.5 \times 10^{-5}$ | $2.6 \times 10^{-3}$ | $6.7 \times 10^{-1}$ | ... |
| **EG13699** | $9.8 \times 10^{-3}$ | $6.4 \times 10^{-1}$ | $5.7 \times 10^{-5}$ | $5.7 \times 10^{-3}$ | $7.8 \times 10^{-1}$ | ... |
| ... | ... | ... | ... | ... | ... | ... |
| ... | ... | ... | ... | ... | ... | ... |
| ... | ... | ... | ... | ... | ... | ... |

Figure S.8: Representation of the p-values matrix. Each column correspond to a pairwise comparison between two datasets. For the sake of readability only 5 columns and 5 rows are reported here. The complete matrix is composed by 36 columns and 2238 rows.

statistical framework, we exploited the False Discovery Rate (FDR) concept and the Benjamini-Hockberg (BH) method correction for multiple testing. In particular, for any given row of the p-values matrix, we set a FDR threshold of 0.01, that is we accepted that the 1% of the possible p-values resulting significant when considered in isolation, might have assumed that value due to randomness. Then, we applied the BH method to adjust the chosen FDR threshold taking into account all the 36 p-values of each single row. Then, we counted in each row how many p-values resulted significant according to the BH method and we defined *reproducible* those Ribo-seq profiles associated with the rows where all the p-values were significant. Following this strategy, we found that out of 2238 possible Ribo-seq profiles only 11 (see Table S.3 and Table 1 in main text) turned out to be reproducible.

To check whether the analysis of the replicates of the considered Samples might yield different results, for each of the replicates contained in the group A (Table S.2) we repeated the process described above. In particular, we performed new iterations of the reproducibility analysis replacing the Sample A1$\alpha$ with its replicate A2$\alpha$, the Sample A3$\beta$ with A4$\beta$ and A5$\beta$, A6$\gamma$ with A7$\gamma$, A12$\theta$ with A13$\theta$ and A14$\iota$ with A15$\iota$, A16$\iota$ and A17$\iota$. It turned out that in all these new 8 cycles of analysis both the number and the type of reproducible Ribo-seq profiles remained unchanged, thus meaning that in the considered cases the choice of the experimental replicates has no influence on reproducibility.

To conclude the reproducibility analysis on the datasets belonging to group A (Table S.2) we checked for the presence of Samples that, in spite of being collected in the

Table S.3: List of the ORFs corresponding to the reproducible Ribo-seq profiles and the p-values associated to each pairwise comparison. This Table is analogous to the p-values matrix depicted in Figure S.8, *i.e.*, the columns contain p-values referring to each performed pairwise comparison.

| **ORF ID** | ***A3β* vs. *A6γ*** | **A3β vs. A10ζ** | **A3β vs. A9ϵ** | **A3β vs. A11η** | A10ζ vs. A9ϵ | A10ζ vs. A11η | A8δ vs. A6γ | A8δ vs. A3β | A8δ vs. A10ζ | A8δ vs. A9ϵ | A8δ vs. A11η | A1α vs. A11η |
|---|---|---|---|---|---|---|---|---|---|---|---|---|
| EG10403 | 1e-08 | 4e-11 | 5e-13 | 1e-09 | 3e-04 | 6e-10 | 3e-10 | 3e-14 | 6e-05 | 5e-10 | 2e-07 | 0e+00 |
| EG11036 | 0e+00 | 0e+00 | 0e+00 | 0e+00 | 0e+00 | 0e+00 | 0e+00 | 0e+00 | 0e+00 | 0e+00 | 0e+00 | 0e+00 |
| EG11037 | 0e+00 | 0e+00 | 0e+00 | 0e+00 | 0e+00 | 0e+00 | 0e+00 | 0e+00 | 0e+00 | 0e+00 | 0e+00 | 0e+00 |
| EG11240 | 3e-03 | 4e-03 | 5e-03 | 3e-03 | 5e-03 | 3e-03 | 3e-04 | 2e-04 | 4e-04 | 1e-04 | 1e-03 | 1e-03 |
| EG11474 | 2e-05 | 1e-07 | 7e-05 | 1e-06 | 8e-06 | 1e-09 | 2e-07 | 7e-06 | 6e-08 | 6e-07 | 4e-07 | 4e-06 |
| EG11773 | 9e-04 | 4e-03 | 4e-03 | 4e-05 | 2e-03 | 3e-03 | 3e-07 | 1e-03 | 3e-03 | 1e-03 | 8e-07 | 3e-11 |
| EG11982 | 3e-06 | 2e-04 | 4e-05 | 9e-08 | 1e-04 | 5e-04 | 7e-06 | 1e-07 | 2e-03 | 1e-06 | 2e-12 | 0e+00 |
| EG11983 | 5e-05 | 5e-05 | 8e-05 | 3e-05 | 2e-03 | 5e-06 | 9e-07 | 2e-09 | 2e-04 | 3e-05 | 2e-05 | 3e-08 |
| EG12323 | 0e+00 | 1e-16 | 5e-14 | 0e+00 | 3e-13 | 3e-16 | 0e+00 | 0e+00 | 0e+00 | 0e+00 | 0e+00 | 0e+00 |
| EG13935 | 1e-06 | 4e-06 | 1e-04 | 3e-07 | 2e-04 | 1e-04 | 1e-06 | 2e-05 | 1e-07 | 4e-08 | 6e-05 | 0e+00 |
| EG14133 | 5e-06 | 1e-06 | 4e-04 | 1e-07 | 5e-05 | 4e-09 | 1e-10 | 6e-06 | 1e-06 | 1e-07 | 8e-09 | 0e+00 |

| **ORF ID** | A1α vs. A3β | A1α vs. A6γ | A1α vs. A8δ | A1α vs. A9ϵ | A1α vs. A10ζ | A6δ vs. A9ϵ | A6δ vs. A10ζ | A6δ vs. A11η | A9ϵ vs. A11η | A12θ vs. A1α | A12θ vs. A3β | A12θ vs. A6γ |
|---|---|---|---|---|---|---|---|---|---|---|---|---|
| EG10403 | 0e+00 | 0e+00 | 0e+00 | 0e+00 | 0e+00 | 4e-09 | 6e-08 | 6e-09 | 0e+00 | 1.6e-10 | 2.8e-10 | 8.1e-11 |
| EG11036 | 0e+00 | 0e+00 | 0e+00 | 0e+00 | 0e+00 | 0e+00 | 0e+00 | 0e+00 | 0e+00 | 0e+00 | 0e+00 | 0e+00 |
| EG11037 | 0e+00 | 0e+00 | 0e+00 | 0e+00 | 0e+00 | 0e+00 | 0e+00 | 0e+00 | 0e+00 | 0e+00 | 0e+00 | 0e+00 |
| EG11240 | 1e-03 | 4e-04 | 3e-04 | 1e-03 | 1e-03 | 2e-03 | 2e-03 | 2e-03 | 8e-04 | 4.9e-04 | 4.2e-04 | 5.8e-04 |
| EG11474 | 9e-06 | 3e-09 | 1e-06 | 5e-05 | 1e-07 | 2e-05 | 2e-07 | 2e-05 | 2e-05 | 2.2e-07 | 9.4e-08 | 3.6e-08 |
| EG11773 | 7e-14 | 0e+00 | 1e-16 | 4e-14 | 0e+00 | 5e-03 | 1e-03 | 8e-06 | 6e-06 | 3.5e-05 | 2.0e-05 | 8.5e-06 |
| EG11982 | 0e+00 | 0e+00 | 0e+00 | 0e+00 | 0e+00 | 1e-06 | 1e-05 | 4e-05 | 1e-06 | 6.1e-07 | 9.4e-07 | 8.1e-07 |
| EG11983 | 4e-15 | 9e-16 | 0e+00 | 9e-15 | 1e-08 | 7e-03 | 1e-03 | 4e-03 | 8e-09 | 1.2e-06 | 8.4e-06 | 3.0e-06 |
| EG12323 | 0e+00 | 0e+00 | 0e+00 | 9e-06 | 2e-09 | 1e-13 | 2e-16 | 0e+00 | 1e-15 | 0.0e+00 | 0.0e+00 | 0.0e+00 |
| EG13935 | 0e+00 | 0e+00 | 0e+00 | 0e+00 | 0e+00 | 6e-05 | 5e-06 | 3e-06 | 2e-07 | 5.7e-07 | 5.6e-07 | 2.0e-07 |
| EG14133 | 0e+00 | 0e+00 | 4e-16 | 9e-14 | 0e+00 | 1e-06 | 3e-12 | 1e-14 | 1e-07 | 5.8e-10 | 5.5e-09 | 1.9e-09 |

| **ORF ID** | A12θ vs. A8δ | A12θ vs. A9ϵ | A12θ vs. A10ζ | A12θ vs. A11η | A12θ vs. A14ι | A14ι vs. A1α | A14ι vs. A3β | A14ι vs. A6γ | A14ι vs. A8δ | A14ι vs. A9ϵ | A14ι vs. A10ζ | A14ι vs. A11η |
|---|---|---|---|---|---|---|---|---|---|---|---|---|
| EG10403 | 1.9e-10 | 2.5e-10 | 2.0e-10 | 1.4e-10 | 2.7e-10 | 2.2e-10 | 2.4e-10 | 1.9e-11 | 3.2e-12 | 2.3e-10 | 6.3e-11 | 1.2e-10 |
| EG11036 | 0e+00 | 0e+00 | 0e+00 | 0e+00 | 0e+00 | 0e+00 | 0e+00 | 0e+00 | 0e+00 | 0e+00 | 0e+00 | 0e+00 |
| EG11037 | 0e+00 | 0e+00 | 0e+00 | 0e+00 | 0e+00 | 0e+00 | 0e+00 | 0e+00 | 0e+00 | 0e+00 | 0e+00 | 0e+00 |
| EG11240 | 5.5e-04 | 1.0e-04 | 2.3e-04 | 3.7e-04 | 4.2e-04 | 8.4e-04 | 3.1e-04 | 7.9e-04 | 6.6e-04 | 7.0e-04 | 8.2e-04 | 7.1e-04 |
| EG11474 | 5.1e-07 | 2.4e-07 | 7.6e-07 | 4.5e-07 | 7.3e-07 | 9.9e-07 | 4.9e-07 | 6.9e-07 | 3.0e-07 | 6.5e-07 | 6.7e-07 | 3.0e-07 |
| EG11773 | 2.0e-05 | 2.1e-05 | 5.4e-06 | 1.6e-05 | 1.3e-05 | 2.5e-05 | 1.8e-05 | 1.9e-05 | 1.7e-05 | 2.4e-06 | 3.0e-06 | 2.0e-05 |
| EG11982 | 3.1e-07 | 3.8e-07 | 4.2e-07 | 2.3e-07 | 9.8e-07 | 8.1e-07 | 7.5e-07 | 8.5e-07 | 1.8e-07 | 7.2e-07 | 8.9e-07 | 9.0e-07 |
| EG11983 | 1.2e-05 | 9.2e-06 | 1.9e-05 | 3.2e-07 | 1.2e-05 | 4.0e-06 | 9.8e-06 | 1.3e-06 | 1.5e-05 | 1.2e-05 | 1.9e-05 | 4.0e-06 |
| EG12323 | 0.0e+00 | 0.0e+00 | 0.0e+00 | 0.0e+00 | 0.0e+00 | 0.0e+00 | 0.0e+00 | 0.0e+00 | 0.0e+00 | 0.0e+00 | 0.0e+00 | 0.0e+00 |
| EG13935 | 7.0e-07 | 6.4e-07 | 4.2e-07 | 8.7e-07 | 5.7e-08 | 6.7e-07 | 1.1e-07 | 1.5e-07 | 6.1e-07 | 1.7e-07 | 2.0e-07 | 3.1e-07 |
| EG14133 | 3.0e-10 | 3.2e-09 | 1.1e-09 | 7.5e-09 | 7.1e-09 | 2.5e-09 | 6.5e-09 | 6.4e-09 | 6.3e-09 | 5.9e-09 | 3.3e-09 | 6.0e-10 |

experimental condition characterising the group, might be affected by unpredictable biases thus influencing the reproducibility results. To do this, we relied on a tailored jackknife approach. Indeed, we considered the p-values matrix and we repeated 9 times the last step of the reproducibility analysis (namely, the identification of the reproducible Ribo-seq profiles) excluding each time one of the datasets of the group A. More in details, for each of the 9 iterations, we removed from the p-values table all the columns referring to comparisons involving the excluded Sample, obtaining a new p-values table composed by 28 - $\binom{8}{2}$ - columns instead of 36. Then, for each row of the new p-values matrix we sampled randomly 8 p-values and we replaced with them the 8 columns removed before. We, then, obtained a 2238 $\times$ 36 *test p-value matrix* in which 8 of the 36 columns were obtained through random sampling. Subsequently, repeating the process used for the original p-values matrix, we tested each row of the 9 test p-value matrices through the FDR - BH method and we counted the number of reproducible Ribo-seq profiles.

After this analysis it turned out that in all the cases in which the Sample A10$\zeta$ was excluded, the number of reproducible Ribo-seq profiles raised from 11 to 49 (see Table S.4 and Table S.5). We interpreted this result as originating from a peculiarity of the sample A10$\zeta$ and, thus, we decided to exclude this Sample from the subsequent analysis.

##### S.2.2 The impact of E. coli's genotype and culture medium on reproducibility

We used our method to evaluate whether performing Ribo-seq using *E. coli* strains or culture media different from those characterising the group A, might affect experimental reproducibility and, thus, the amount of high-resolution ribosome profiles. To do this, we examined all the samples belonging to the groups B, C, D, E, F, G listed in Table S.2 and we compared them with the Samples of group A which represent our benchmark.

###### S.2.2.1 Analysing the reproducibility of Ribo-seq profiles for different culture media

We started this comparative analysis considering group B at first which refers to Ribo-seq data collected from *E. coli* K-12 MG1655 (the same genotype chosen in group A) cultured in the LB medium. Thus, comparing Samples belonging to group B with those present in group A allows us to evaluate whether the experimental variable culture medium (in this case LB *vs.* MOPS-rich medium) might influence the reproducibility of Ribo-seq. More in detail, we considered the Samples B1$\alpha$ and B3$\beta$ at first and, for the both of them, we generated the Ribo-seq profiles corresponding to the 2238 ORFs

Table S.4: List of the ORFs corresponding to the reproducible Ribo-seq profiles, once the Sample A10$\zeta$ was not considered in the analysis. This Table is analogous to the p-values matrix depicted in Figure S.8, *i.e.*, the columns contain p-values referring to each performed pairwise comparison. The entries in common with Table S.2 are italicised.

| **ORF ID** | ***A3$\beta$* vs. *A6$\gamma$*** | **A3$\beta$ vs. A9$\epsilon$** | **A3$\beta$ vs. A11$\eta$** | A8$\delta$ vs. A6$\gamma$ | A8$\delta$ vs. A3$\beta$ | A8$\delta$ vs. A9$\epsilon$ | A8$\delta$ vs. A11$\eta$ | A1$\alpha$ vs. A11$\eta$ | A1$\alpha$ vs. A3$\beta$ | A1$\alpha$ vs. A6$\gamma$ | A1$\alpha$ vs. A8$\delta$ | A1$\alpha$ vs. A9$\epsilon$ | A6$\delta$ vs. A9$\epsilon$ | A6$\delta$ vs. A11$\eta$ | A9$\epsilon$ vs. A11$\eta$ | A12$\theta$ vs. A1$\alpha$ | A12$\theta$ vs. A3$\beta$ | A12$\theta$ vs. A6$\gamma$ | A12$\theta$ vs. A8$\delta$ | A12$\theta$ vs. A9$\epsilon$ | A12$\theta$ vs. A11$\eta$ | A12$\theta$ vs. A14$\iota$ | A14$\iota$ vs. A1$\alpha$ | A14$\iota$ vs. A3$\beta$ | A14$\iota$ vs. A6$\gamma$ | A14$\iota$ vs. A8$\delta$ | A14$\iota$ vs. A9$\epsilon$ | A14$\iota$ vs. A11$\eta$ |
|---|---|---|---|---|---|---|---|---|---|---|---|---|---|---|---|---|---|---|---|---|---|---|---|---|---|---|---|---|
| EG10015 | 9e-04 | 6e-04 | 3e-07 | 3e-05 | 1e-04 | 2e-05 | 2e-05 | 9e-08 | 1e-05 | 9e-04 | 5e-03 | 1e-03 | 2e-03 | 1e-05 | 2e-07 | 1.5e-05 | 1.0e-05 | 1.2e-05 | 2.9e-06 | 1.0e-05 | 4.8e-06 | 9.7e-06 | 4.6e-06 | 4.8e-06 | 6.5e-06 | 1.3e-05 | 1.1e-05 | 4.6e-06 |
| EG10017 | 2e-04 | 1e-03 | 2e-03 | 2e-05 | 7e-05 | 9e-04 | 4e-05 | 1e-03 | 1e-05 | 3e-08 | 1e-06 | 3e-03 | 1e-03 | 6e-05 | 2e-05 | 4.5e-05 | 4.5e-06 | 2.7e-07 | 5.6e-05 | 2.6e-05 | 1.3e-05 | 8.9e-06 | 8.5e-06 | 5.0e-05 | 3.1e-05 | 2.1e-05 | 2.3e-05 | 1.4e-05 |
| EG10024 | 6e-07 | 8e-07 | 2e-04 | 3e-11 | 5e-12 | 1e-15 | 7e-10 | 0e+00 | 0e+00 | 0e+00 | 0e+00 | 0e+00 | 3e-11 | 4e-11 | 7e-09 | 1.0e-12 | 2.7e-12 | 2.5e-12 | 3.0e-12 | 1.8e-12 | 2.4e-12 | 1.9e-12 | 2.8e-12 | 2.4e-12 | 1.7e-12 | 6.1e-13 | 3.4e-12 | 1.5e-12 |
| EG10034 | 1e-03 | 9e-03 | 2e-05 | 9e-12 | 4e-05 | 8e-10 | 1e-10 | 0e+00 | 0e+00 | 0e+00 | 0e+00 | 0e+00 | 1e-07 | 1e-09 | 2e-07 | 4.1e-11 | 2.8e-11 | 2.9e-11 | 5.6e-11 | 4.9e-11 | 1.1e-11 | 8.7e-11 | 8.5e-11 | 1.8e-11 | 3.7e-11 | 2.1e-11 | 5.4e-11 | 9.8e-11 |
| EG10156 | 4e-03 | 2e-03 | 6e-04 | 2e-04 | 1e-05 | 6e-07 | 4e-05 | 8e-08 | 2e-05 | 2e-04 | 1e-05 | 1e-03 | 4e-03 | 8e-03 | 7e-09 | 2.8e-05 | 3.2e-05 | 1.9e-06 | 1.1e-05 | 3.0e-05 | 1.2e-05 | 5.4e-06 | 1.7e-05 | 1.5e-05 | 2.8e-06 | 2.1e-05 | 5.3e-06 | 3.5e-05 |
| EG10178 | 1e-03 | 8e-04 | 2e-05 | 3e-06 | 2e-05 | 7e-05 | 1e-04 | 2e-13 | 2e-10 | 6e-09 | 3e-10 | 6e-08 | 2e-05 | 3e-05 | 8e-06 | 5.4e-07 | 8.5e-07 | 4.4e-06 | 6.1e-06 | 1.2e-06 | 2.2e-06 | 4.6e-06 | 4.5e-06 | 2.3e-06 | 1.4e-06 | 9.8e-07 | 3.6e-06 | 3.5e-06 |
| EG10215 | 9e-03 | 6e-04 | 1e-02 | 3e-03 | 2e-03 | 8e-06 | 3e-09 | 0e+00 | 0e+00 | 0e+00 | 0e+00 | 0e+00 | 4e-04 | 8e-04 | 6e-07 | 4.0e-07 | 2.3e-07 | 4.8e-07 | 4.7e-08 | 3.4e-07 | 1.7e-08 | 3.3e-07 | 5.1e-08 | 5.6e-07 | 4.5e-07 | 2.9e-07 | 9.0e-08 | 5.6e-07 |
| EG10231 | 6e-03 | 6e-05 | 3e-04 | 2e-11 | 4e-04 | 2e-10 | 1e-10 | 0e+00 | 0e+00 | 0e+00 | 0e+00 | 0e+00 | 2e-05 | 4e-08 | 6e-08 | 3.6e-12 | 4.1e-11 | 7.4e-11 | 9.8e-11 | 4.6e-11 | 9.8e-11 | 1.2e-11 | 5.8e-11 | 1.2e-11 | 1.6e-11 | 7.3e-11 | 5.5e-11 | 3.3e-11 |
| EG10245 | 8e-05 | 7e-09 | 2e-07 | 2e-08 | 5e-04 | 2e-06 | 1e-09 | 0e+00 | 0e+00 | 0e+00 | 0e+00 | 0e+00 | 2e-08 | 2e-08 | 1e-11 | 1.1e-10 | 5.0e-10 | 4.8e-10 | 6.5e-11 | 8.9e-10 | 6.7e-10 | 8.4e-10 | 2.9e-10 | 5.5e-10 | 7.3e-10 | 3.4e-10 | 6.3e-10 | 1.1e-10 |
| EG10276 | 3e-04 | 9e-03 | 3e-04 | 4e-07 | 7e-04 | 1e-06 | 2e-03 | 0e+00 | 0e+00 | 0e+00 | 0e+00 | 0e+00 | 3e-03 | 3e-05 | 1e-04 | 1.7e-07 | 8.8e-08 | 1.4e-07 | 1.2e-07 | 3.8e-07 | 8.1e-07 | 4.8e-08 | 2.4e-07 | 5.9e-07 | 3.8e-07 | 3.4e-07 | 5.0e-07 | 7.0e-07 |
| *EG10403* | 1e-08 | 5e-13 | 1e-09 | 3e-10 | 3e-14 | 5e-10 | 2e-07 | 0e+00 | 0e+00 | 0e+00 | 0e+00 | 0e+00 | 4e-09 | 6e-09 | 0e+00 | 1.6e-14 | 7.1e-15 | 4.3e-15 | 7.8e-15 | 2.9e-14 | 1.7e-14 | 2.9e-15 | 4.4e-15 | 3.7e-15 | 2.9e-14 | 2.1e-14 | 6.7e-15 | 5.4e-15 |
| EG10424 | 6e-03 | 4e-05 | 1e-04 | 4e-07 | 1e-09 | 1e-05 | 5e-08 | 0e+00 | 0e+00 | 4e-16 | 0e+00 | 0e+00 | 4e-06 | 8e-08 | 2e-13 | 8.9e-10 | 8.7e-10 | 1.9e-10 | 5.8e-10 | 6.3e-10 | 8.8e-10 | 7.5e-10 | 8.7e-10 | 7.4e-10 | 3.8e-10 | 8.1e-11 | 5.2e-10 | 6.8e-10 |
| EG10492 | 5e-04 | 1e-02 | 1e-05 | 1e-07 | 3e-05 | 2e-09 | 6e-10 | 0e+00 | 0e+00 | 0e+00 | 0e+00 | 0e+00 | 1e-04 | 3e-05 | 9e-07 | 1.6e-09 | 6.3e-10 | 1.0e-09 | 3.3e-10 | 1.2e-09 | 1.0e-09 | 1.5e-09 | 3.7e-10 | 1.9e-09 | 3.0e-10 | 7.6e-10 | 1.1e-09 | 9.3e-10 |
| EG10505 | 5e-05 | 2e-03 | 4e-04 | 1e-11 | 5e-06 | 3e-15 | 1e-08 | 0e+00 | 0e+00 | 0e+00 | 0e+00 | 0e+00 | 5e-05 | 4e-06 | 4e-08 | 9.4e-12 | 8.7e-12 | 9.8e-12 | 1.0e-12 | 1.1e-12 | 7.4e-12 | 5.7e-12 | 6.2e-12 | 7.2e-12 | 1.0e-11 | 7.5e-12 | 2.7e-12 | 3.0e-13 |
| EG10534 | 1e-03 | 6e-04 | 8e-05 | 3e-05 | 3e-07 | 3e-08 | 5e-07 | 0e+00 | 0e+00 | 0e+00 | 0e+00 | 0e+00 | 2e-06 | 9e-07 | 1e-10 | 1.3e-08 | 4.2e-09 | 3.2e-10 | 1.6e-08 | 2.8e-08 | 2.5e-08 | 1.2e-08 | 7.7e-09 | 2.5e-09 | 2.1e-08 | 1.9e-08 | 2.4e-08 | 5.9e-09 |
| EG10542 | 3e-03 | 2e-05 | 5e-05 | 1e-11 | 4e-04 | 3e-10 | 2e-13 | 6e-11 | 1e-10 | 3e-15 | 1e-16 | 3e-14 | 5e-06 | 5e-04 | 8e-15 | 5.9e-11 | 6.0e-11 | 2.7e-11 | 3.5e-11 | 2.2e-11 | 3.1e-11 | 3.5e-11 | 4.1e-11 | 4.9e-11 | 4.2e-11 | 4.7e-12 | 2.4e-11 | 5.6e-11 |
| EG10618 | 3e-04 | 2e-03 | 1e-08 | 2e-11 | 3e-06 | 9e-12 | 2e-09 | 0e+00 | 0e+00 | 0e+00 | 0e+00 | 0e+00 | 5e-05 | 7e-08 | 6e-06 | 1.1e-11 | 1.3e-11 | 2.5e-12 | 9.6e-12 | 1.7e-11 | 1.8e-11 | 5.7e-12 | 5.0e-12 | 5.9e-12 | 8.4e-12 | 1.4e-11 | 1.0e-11 | 1.2e-11 |
| EG10635 | 3e-03 | 9e-04 | 7e-03 | 1e-05 | 2e-06 | 3e-07 | 2e-05 | 0e+00 | 0e+00 | 0e+00 | 0e+00 | 0e+00 | 3e-03 | 2e-04 | 3e-04 | 2.4e-07 | 1.7e-06 | 1.9e-06 | 3.1e-07 | 4.5e-07 | 9.6e-07 | 6.8e-07 | 5.8e-07 | 1.5e-06 | 4.6e-07 | 7.3e-07 | 8.6e-07 | 1.1e-06 |
| EG10670 | 9e-05 | 6e-03 | 1e-03 | 2e-04 | 8e-07 | 3e-03 | 7e-06 | 4e-04 | 2e-03 | 5e-05 | 1e-03 | 5e-05 | 8e-04 | 7e-07 | 3e-03 | 1.3e-04 | 1.8e-04 | 1.5e-04 | 5.8e-06 | 1.5e-05 | 1.3e-04 | 2.0e-04 | 1.1e-04 | 7.2e-05 | 1.6e-04 | 1.3e-04 | 3.5e-05 | 7.6e-05 |
| EG10686 | 4e-03 | 1e-03 | 3e-03 | 6e-08 | 2e-04 | 5e-04 | 6e-07 | 0e+00 | 0e+00 | 0e+00 | 0e+00 | 0e+00 | 2e-03 | 9e-06 | 7e-05 | 5.5e-07 | 2.4e-07 | 4.3e-07 | 4.5e-07 | 2.1e-07 | 2.9e-07 | 3.7e-07 | 1.9e-08 | 3.5e-07 | 5.4e-07 | 2.5e-07 | 5.5e-07 | 4.1e-07 |
| EG10696 | 1e-07 | 2e-04 | 2e-05 | 4e-09 | 6e-09 | 1e-10 | 3e-10 | 0e+00 | 0e+00 | 0e+00 | 0e+00 | 0e+00 | 3e-05 | 3e-07 | 9e-10 | 3.0e-11 | 2.1e-10 | 2.2e-10 | 3.1e-11 | 1.1e-10 | 7.7e-11 | 7.5e-11 | 1.5e-10 | 1.8e-11 | 1.8e-10 | 1.6e-10 | 8.0e-11 | 6.9e-11 |
| EG10766 | 4e-04 | 6e-04 | 6e-05 | 3e-05 | 9e-08 | 2e-05 | 4e-09 | 1e-07 | 2e-03 | 5e-03 | 4e-04 | 3e-05 | 2e-04 | 3e-06 | 8e-09 | 1.4e-05 | 1.8e-05 | 2.4e-05 | 2.9e-05 | 6.8e-06 | 1.9e-05 | 3.9e-06 | 1.1e-05 | 2.3e-05 | 5.8e-06 | 2.9e-05 | 3.0e-06 | 1.1e-06 |
| EG10781 | 1e-08 | 2e-06 | 2e-05 | 6e-09 | 7e-08 | 3e-05 | 4e-10 | 0e+00 | 0e+00 | 0e+00 | 0e+00 | 0e+00 | 2e-06 | 4e-07 | 3e-05 | 6.4e-10 | 2.2e-09 | 3.3e-09 | 5.3e-09 | 1.6e-09 | 1.9e-09 | 5.4e-09 | 4.0e-09 | 4.7e-09 | 5.2e-09 | 2.3e-09 | 4.6e-09 | 5.5e-09 |
| EG10859 | 3e-05 | 1e-04 | 2e-13 | 1e-10 | 9e-11 | 3e-11 | 2e-05 | 0e+00 | 0e+00 | 0e+00 | 0e+00 | 0e+00 | 7e-05 | 1e-06 | 4e-07 | 9.3e-12 | 2.0e-11 | 1.7e-11 | 2.5e-12 | 1.2e-11 | 2.4e-11 | 1.7e-11 | 1.8e-11 | 8.1e-12 | 1.7e-11 | 2.3e-11 | 1.4e-11 | 1.1e-11 |
| EG10894 | 2e-04 | 3e-05 | 9e-04 | 4e-12 | 6e-10 | 7e-16 | 6e-15 | 0e+00 | 0e+00 | 0e+00 | 0e+00 | 0e+00 | 3e-09 | 2e-06 | 2e-12 | 4.9e-17 | 1.6e-15 | 3.4e-15 | 1.6e-16 | 4.7e-15 | 4.1e-16 | 5.9e-15 | 3.5e-15 | 3.8e-15 | 3.5e-15 | 8.6e-17 | 4.8e-15 | 4.6e-15 |
| EG10921 | 5e-04 | 2e-03 | 3e-04 | 7e-06 | 2e-05 | 2e-04 | 5e-04 | 0e+00 | 0e+00 | 0e+00 | 0e+00 | 0e+00 | 2e-03 | 2e-03 | 7e-05 | 1.2e-05 | 1.0e-05 | 1.8e-05 | 6.1e-06 | 7.6e-06 | 1.9e-05 | 8.3e-06 | 3.8e-06 | 2.6e-06 | 1.6e-05 | 1.1e-05 | 9.9e-06 | 6.0e-06 |
| EG10936 | 1e-02 | 6e-05 | 2e-03 | 8e-06 | 1e-10 | 1e-09 | 4e-07 | 0e+00 | 0e+00 | 0e+00 | 0e+00 | 0e+00 | 1e-05 | 3e-05 | 2e-09 | 1.2e-10 | 4.2e-10 | 2.3e-10 | 3.9e-10 | 3.2e-10 | 3.6e-10 | 4.5e-10 | 2.1e-10 | 4.1e-10 | 8.4e-10 | 4.3e-10 | 6.3e-10 | 8.2e-11 |
| EG10979 | 6e-03 | 8e-05 | 6e-07 | 5e-09 | 1e-11 | 3e-06 | 2e-10 | 0e+00 | 0e+00 | 0e+00 | 0e+00 | 0e+00 | 3e-05 | 2e-06 | 1e-07 | 1.0e-10 | 1.7e-10 | 1.5e-10 | 1.8e-11 | 1.8e-13 | 1.3e-10 | 1.4e-10 | 1.2e-10 | 1.5e-10 | 1.0e-10 | 1.4e-10 | 1.4e-10 | 1.9e-10 |
| EG10998 | 2e-04 | 4e-04 | 5e-09 | 9e-06 | 2e-07 | 7e-09 | 2e-12 | 0e+00 | 0e+00 | 0e+00 | 0e+00 | 0e+00 | 2e-04 | 4e-06 | 2e-10 | 4.8e-11 | 1.4e-10 | 9.3e-11 | 1.6e-10 | 7.7e-11 | 6.6e-11 | 1.5e-10 | 1.8e-10 | 1.3e-10 | 1.6e-10 | 6.5e-11 | 3.3e-11 | 1.7e-10 |
| EG11036 | 0e+00 | 0e+00 | 0e+00 | 0e+00 | 0e+00 | 0e+00 | 0e+00 | 0e+00 | 0e+00 | 0e+00 | 0e+00 | 0e+00 | 0e+00 | 0e+00 | 0e+00 | 0e+00 | 0e+00 | 0e+00 | 0e+00 | 0e+00 | 0e+00 | 0e+00 | 0e+00 | 0e+00 | 0e+00 | 0e+00 | 0e+00 | 0e+00 |
| *EG11037* | 0e+00 | 0e+00 | 0e+00 | 0e+00 | 0e+00 | 0e+00 | 0e+00 | 0e+00 | 0e+00 | 0e+00 | 0e+00 | 0e+00 | 0e+00 | 0e+00 | 0e+00 | 0e+00 | 0e+00 | 0e+00 | 0e+00 | 0e+00 | 0e+00 | 0e+00 | 0e+00 | 0e+00 | 0e+00 | 0e+00 | 0e+00 | 0e+00 |
| *EG11240* | 3e-03 | 5e-03 | 3e-03 | 3e-04 | 2e-04 | 1e-04 | 1e-03 | 1e-03 | 1e-03 | 4e-04 | 3e-04 | 1e-03 | 2e-03 | 2e-03 | 8e-04 | 6.3e-04 | 5.3e-04 | 4.7e-04 | 7.0e-04 | 9.2e-04 | 3.1e-04 | 4.8e-04 | 4.7e-04 | 9.6e-04 | 7.8e-04 | 9.2e-04 | 8.1e-04 | 6.4e-04 |
| EG11259 | 7e-07 | 3e-05 | 3e-04 | 7e-07 | 4e-05 | 3e-04 | 2e-03 | 0e+00 | 0e+00 | 0e+00 | 0e+00 | 0e+00 | 2e-03 | 1e-04 | 1e-04 | 6.0e-07 | 6.4e-07 | 3.1e-07 | 6.8e-07 | 5.3e-07 | 1.6e-07 | 3.7e-07 | 9.7e-08 | 1.5e-07 | 3.4e-07 | 2.9e-07 | 6.0e-08 | 5.2e-07 |
| EG11352 | 2e-04 | 4e-03 | 2e-04 | 7e-08 | 2e-05 | 2e-06 | 4e-10 | 0e+00 | 0e+00 | 0e+00 | 0e+00 | 0e+00 | 1e-03 | 4e-07 | 2e-04 | 6.0e-08 | 5.2e-08 | 6.2e-08 | 2.3e-08 | 4.1e-08 | 1.3e-08 | 6.9e-08 | 6.0e-08 | 3.3e-08 | 4.0e-08 | 1.6e-08 | 3.1e-08 | 5.7e-08 |
| EG11370 | 4e-03 | 8e-03 | 1e-04 | 5e-05 | 3e-04 | 2e-09 | 7e-07 | 0e+00 | 0e+00 | 0e+00 | 0e+00 | 0e+00 | 2e-03 | 4e-05 | 1e-06 | 1.6e-07 | 6.4e-07 | 3.0e-07 | 4.7e-07 | 1.7e-07 | 1.5e-07 | 6.7e-08 | 2.3e-07 | 1.1e-07 | 6.4e-07 | 6.0e-07 | 6.5e-07 | 3.6e-07 |
| EG11459 | 4e-03 | 2e-05 | 3e-03 | 6e-03 | 3e-04 | 5e-06 | 1e-04 | 0e+00 | 5e-12 | 9e-16 | 4e-10 | 5e-10 | 1e-03 | 4e-08 | 1e-06 | 5.2e-07 | 7.4e-07 | 6.8e-07 | 4.8e-07 | 5.4e-07 | 7.4e-07 | 6.3e-07 | 1.2e-07 | 2.7e-07 | 9.5e-07 | 5.9e-07 | 5.1e-07 | 1.5e-07 |
| *EG11474* | 2e-05 | 7e-05 | 1e-06 | 2e-07 | 7e-06 | 6e-07 | 4e-07 | 4e-06 | 9e-06 | 3e-09 | 1e-06 | 5e-05 | 2e-05 | 2e-05 | 2e-05 | 4.0e-06 | 1.8e-06 | 3.0e-06 | 2.0e-06 | 2.5e-06 | 7.2e-07 | 3.2e-06 | 3.0e-06 | 3.5e-07 | 2.6e-06 | 2.5e-06 | 3.0e-06 | 2.5e-06 |
| EG11569 | 3e-03 | 2e-03 | 1e-04 | 1e-08 | 8e-06 | 5e-10 | 1e-10 | 0e+00 | 0e+00 | 0e+00 | 0e+00 | 0e+00 | 2e-04 | 2e-09 | 3e-06 | 4.5e-10 | 2.0e-11 | 3.0e-11 | 2.6e-10 | 1.7e-10 | 4.4e-10 | 9.4e-11 | 8.8e-11 | 3.1e-10 | 4.7e-12 | 6.4e-11 | 4.4e-10 | 2.2e-10 |
| EG11612 | 2e-04 | 4e-07 | 2e-10 | 4e-03 | 7e-04 | 3e-05 | 4e-06 | 0e+00 | 0e+00 | 0e+00 | 0e+00 | 0e+00 | 3e-04 | 1e-03 | 5e-07 | 9.0e-08 | 1.6e-07 | 2.9e-07 | 1.8e-07 | 3.9e-07 | 1.5e-07 | 6.0e-08 | 3.9e-07 | 2.2e-07 | 3.6e-07 | 1.2e-07 | 1.8e-09 | 5.8e-08 |
| *EG11773* | 9e-04 | 4e-03 | 4e-05 | 3e-07 | 1e-03 | 1e-03 | 8e-07 | 3e-11 | 7e-14 | 0e+00 | 1e-16 | 4e-14 | 5e-03 | 8e-06 | 6e-06 | 1.4e-07 | 1.4e-07 | 3.0e-08 | 1.3e-07 | 4.1e-07 | 4.8e-07 | 1.7e-07 | 6.4e-07 | 4.6e-07 | 2.3e-08 | 3.8e-07 | 5.7e-07 | 6.7e-07 |
| *EG11982* | 3e-06 | 4e-05 | 9e-08 | 7e-06 | 1e-07 | 1e-06 | 2e-12 | 0e+00 | 0e+00 | 0e+00 | 0e+00 | 0e+00 | 1e-06 | 4e-05 | 1e-06 | 3.0e-08 | 7.1e-08 | 3.5e-08 | 3.6e-08 | 5.3e-08 | 6.8e-08 | 8.2e-08 | 6.0e-08 | 7.0e-08 | 5.8e-09 | 6.7e-08 | 8.8e-08 | 7.4e-08 |
| *EG11983* | 5e-05 | 8e-05 | 3e-05 | 9e-07 | 2e-09 | 3e-05 | 2e-05 | 3e-08 | 4e-15 | 9e-16 | 0e+00 | 9e-15 | 7e-03 | 4e-03 | 8e-09 | 1.8e-09 | 3.7e-09 | 2.3e-08 | 1.4e-08 | 2.6e-08 | 2.8e-09 | 3.7e-09 | 8.2e-09 | 1.5e-08 | 5.8e-09 | 5.8e-09 | 7.1e-09 | 1.0e-08 |
| *EG12323* | 0e+00 | 5e-14 | 0e+00 | 0e+00 | 0e+00 | 0e+00 | 0e+00 | 0e+00 | 0e+00 | 0e+00 | 0e+00 | 9e-06 | 1e-13 | 0e+00 | 1e-15 | 0.0e+00 | 0.0e+00 | 0.0e+00 | 0.0e+00 | 0.0e+00 | 0.0e+00 | 0.0e+00 | 0.0e+00 | 0.0e+00 | 0.0e+00 | 0.0e+00 | 0.0e+00 | 0.0e+00 |
| EG13134 | 2e-03 | 7e-03 | 3e-04 | 3e-05 | 3e-06 | 5e-06 | 3e-06 | 0e+00 | 0e+00 | 0e+00 | 0e+00 | 0e+00 | 5e-04 | 5e-07 | 2e-05 | 1.9e-06 | 2.6e-06 | 2.5e-06 | 2.2e-06 | 2.6e-06 | 1.1e-06 | 7.5e-07 | 2.0e-06 | 3.8e-07 | 8.4e-07 | 2.6e-06 | 1.9e-06 | 1.1e-06 |
| EG13183 | 2e-06 | 1e-02 | 8e-08 | 4e-06 | 7e-04 | 6e-07 | 1e-04 | 0e+00 | 0e+00 | 0e+00 | 0e+00 | 0e+00 | 1e-04 | 1e-15 | 1e-04 | 1.1e-08 | 2.1e-08 | 5.8e-08 | 7.3e-08 | 2.4e-08 | 4.1e-08 | 7.7e-08 | 7.0e-08 | 4.8e-09 | 4.0e-08 | 6.2e-08 | 4.4e-09 | 2.0e-08 |
| EG13612 | 3e-04 | 3e-04 | 1e-05 | 2e-05 | 5e-06 | 7e-04 | 3e-04 | 0e+00 | 0e+00 | 0e+00 | 0e+00 | 0e+00 | 6e-03 | 2e-03 | 2e-04 | 7.0e-06 | 3.3e-06 | 7.4e-06 | 8.2e-06 | 3.8e-06 | 6.5e-06 | 4.5e-06 | 8.9e-06 | 6.8e-06 | 7.4e-06 | 5.3e-06 | 9.7e-06 | 4.5e-06 |
| EG13699 | 1e-02 | 6e-05 | 6e-03 | 1e-06 | 2e-04 | 2e-07 | 1e-08 | 5e-11 | 2e-13 | 0e+00 | 3e-09 | 6e-04 | 2e-03 | 4e-05 | 1e-07 | 1.1e-07 | 2.0e-07 | 5.0e-08 | 1.9e-07 | 2.5e-08 | 1.6e-07 | 1.3e-07 | 1.7e-07 | 1.5e-08 | 1.7e-07 | 3.0e-10 | 9.0e-08 | 2.1e-08 |
| *EG13935* | 1e-06 | 1e-04 | 3e-07 | 1e-06 | 2e-05 | 4e-08 | 6e-05 | 0e+00 | 0e+00 | 0e+00 | 0e+00 | 0e+00 | 6e-05 | 3e-06 | 2e-07 | 1.2e-07 | 1.4e-07 | 1.8e-07 | 1.2e-07 | 1.2e-07 | 1.7e-07 | 1.0e-07 | 1.8e-07 | 6.4e-08 | 1.9e-07 | 3.5e-08 | 4.0e-08 | 8.2e-08 |
| *EG14133* | 5e-06 | 4e-04 | 1e-07 | 1e-10 | 6e-06 | 1e-07 | 8e-09 | 0e+00 | 0e+00 | 0e+00 | 4e-16 | 9e-14 | 1e-06 | 1e-14 | 1e-07 | 3.8e-11 | 7.6e-11 | 4.1e-11 | 9.8e-11 | 4.9e-12 | 3.9e-11 | 4.4e-11 | 7.5e-11 | 7.5e-12 | 1.6e-12 | 8.4e-11 | 8.9e-11 | 7.4e-11 |

| Gene | Function | Length (aa) |
|---|---|---|
| gltB | Glutamate synthase, large subunit | 1486 |
| tufA | Translation elongation factor EF-Tu 1 | 394 |
| tufB | Translation elongation factor EF-Tu 2 | 394 |
| hokB | Small toxic membrane polypeptide | 49 |
| ubiJ | Aerobic ubiquinone synthesis protein, SCP2 family protein | 201 |
| nuoM | NADH:ubiquinone oxidoreductase subunit M, complex I | 509 |
| wbbH | O-antigen polymerase | 388 |
| wbbI | d-Galf:alpha-d-Glc beta-1,6-galactofuranosyltransferase | 330 |
| yfaD | Transposase31 family protein, function unknown | 299 |
| rsxC | SoxR iron-sulfur cluster reduction factor component | 740 |
| gtrS | Serotype-specific glucosyl transferase | 443 |
| rodZ | transmembrane component of cytoskeleton | 337 |
| ispB | all-trans-octaprenyl-diphosphate synthase | 323 |
| aceE | pyruvate dehydrogenase | 887 |
| alaS | alanine-tRNA ligase/DNA-binding transcriptional repressor | 876 |
| ClpA | ClpA ATP-dependent protease specificity component | 758 |
| cyoA | cytochrome bo3 ubiquinol oxidase subunit 2 | 315 |
| DeaD | ATP-dependent RNA helicase DeaD | 629 |
| dld | D-lactate dehydrogenase | 571 |
| dnaX | DNA polymerase III subunit gamma | 431 |
| accC | biotin carboxylase | 449 |
| gyrB | DNA gyrase subunit B | 804 |
| ileS | isoleucine-tRNA ligase | 938 |
| infB | translation initiation factor IF-2 | 890 |
| ligA | DNA ligase | 671 |
| lon | Lon protease | 784 |
| MukB | chromosome partitioning protein MukB | 1486 |
| nagE | N-acetylglucosamine specific PTS enzyme II | 648 |
| ompC | outer membrane porin C | 367 |
| parC | dimer of DNA topoisomerase IV subunit A | 752 |
| pepN | aminopeptidase N | 870 |
| secY | Sec translocon subunit SecY | 443 |
| pssA | phosphatidylserine synthase | 451 |
| rne | ribonuclease E | 1061 |
| rpoB | RNA polymerase subunit beta | 1342 |
| ftnA | ferritin iron-storage complex | 165 |
| secA | protein translocation ATPase | 901 |
| sucA | 2-oxoglutarate decarboxylase, thiamine-requiring | 933 |
| thrA | fused aspartate kinase/homoserine dehydrogenase 1 | 820 |
| rnr | RNase R | 813 |
| waaO | UDP-D-glucose:(glucosyl)LPS alpha-1,3-glucosyltransferase | 339 |
| ubiA | 4-hydroxybenzoate octaprenyltransferase | 290 |
| YifK | putative transporter YifK | 461 |
| lptD | lipopolysaccharide assembly protein LptD | 784 |
| proP | osmolyte:H+ symporter ProP | 500 |
| uup | ATP-binding protein | 635 |
| ydbK | putative pyruvate-flavodoxin oxidoreductase | 1174 |
| dxs | 1-deoxy-D-xylulose-5-phosphate synthase | 620 |
| ycaO | ribosomal prot. S12 methylthiotransferase accessory factor YcaO | 586 |

Table S.5: Genes with high resolution Ribo-seq profile, after excluding the dataset GSE85540 analysed previously. Then, according to our method, we generated the corresponding digitalised profiles and we compared them pairwise with the homologous ones that we computed previously for the 8 datasets belonging to group A. Recall that, as discussed in Section S.2.1 we ruled out the sample A$\zeta$ from the 9 samples belonging to group A. These comparisons yielded two 2238 $\times$ 8 scores matrices (one for B1$\alpha$ and one for B3$\beta$) and two 2238 $\times$ 8 null distribution matrices. Mapping the similarity scores on the corresponding null distributions we obtained two 2238 $\times$ 8 p-values matrices (call them *test p-values matrices*) that quantify the degree of similarity between the single Ribo-seq profiles coming from the two Samples of group B and the benchmark profiles. Finally, we composed columnwise the 2238 $\times$ 28 "benchmark" p-values matrix with the test p-values matrices, obtaining two 2238 $\times$ 36 matrices. For each of them, we performed rowwise the FDR-BH test and we counted the amount of rows in which all the p-values resulted significant. It turned out that this condition was verified for 49 rows in both the composed matrices and that the ORFs corresponding to these rows were the same yielded from the analysis of the 8 group A Samples (Table S.4). We obtained similar results also when the replicates of Samples B3$\beta$ and B1$\alpha$ (Samples B4$\beta$, B5$\beta$ and B2$\alpha$ respectively) were considered. We interpreted these results, summarised in Table S.6 observing that the choice of the LB medium instead of the MOPS rich one has no impact on the reproducibility of Ribo-seq, at least for the analysed Samples.

| **Group** | **Tested Samples** | **Culture Medium** | **Tested ORFs** | **Reproducible Profiles** |
|---|---|---|---|---|
| B | B1$\alpha$ | LB | 2238 | 49 |
| | B2$\alpha$ | LB | 2238 | 49 |
| | B3$\beta$ | LB | 2238 | 49 |
| | B4$\beta$ | LB | 2238 | 49 |
| | B5$\beta$ | LB | 2238 | 49 |

Table S.6: Results of the comparative analysis to check the impact on experimental reproducibility when the LB culture medium is used instead of the MOPS rich medium. Column 1: Sample's group; Column 2: Sample's ID; Column 3: culture medium used; Column 4: number of tested ORFs, *i.e.*, number of ORFs in common between the tested Sample and the samples belonging to the benchmark group A; Column 5: number of reproducible profiles identified from the comparative analysis. **Note**: the ORFs ruled out from the comparative analysis because not in common between the tested datasets do not contain any of the reproducible profiles listed in Table S.4.

Following the strategy described above, we tested the Samples of group C against our benchmark (group A). Also in this case, the impact on experimental reproducibility of the variable culture medium was investigated, given that the same *E. coli*'s genotype characterises both groups A and C which differ for the chosen culture medium (Minimal Medium instead of MOPS-rich medium). In this comparative analysis, only the 2214 ORFs in common between the Samples in groups A and C and were considered. Noteworthy, this set includes all the 49 ORFs reported in Table S.4. When the samples C1$\alpha$, C3$\beta$ and the corresponding replicates were challenged against the benchmark, it turned out that only 24 to 26 Ribo-seq profiles exhibited significant reproducibility, as reported in Table S.7. In our perspective, this relevant drop in the number of reproducible Ribo-seq profiles calls for poor consistency between the Ribo-seq experiments conducted growing *E. coli* k-12 MG1655 in a MOPS-based medium or in a minimal medium.

| **Group** | **Tested Samples** | **Growth Medium** | **Tested ORFs** | **Reproducible Profiles** |
|---|---|---|---|---|
| C | C1$\alpha$ | Minimal Medium | 2214 | 24 |
| | C2$\alpha$ | Minimal Medium | 2214 | 24 |
| | C3$\beta$ | Minimal Medium | 2214 | 27 |
| | C4$\beta$ | Minimal Medium | 2214 | 27 |

Table S.7: Results of the comparative analysis to check the impact on experimental reproducibility when the minimal culture medium is used instead of the MOPS rich medium. Column 1: Sample's group; Column 2: Sample's ID; Column 3: culture medium used; Column 4: number of tested ORFs, *i.e.*, number of ORFs in common between the tested Sample and the samples belonging to the benchmark group A; Column 5: number of reproducible profiles identified from the comparative analysis. **Note**: the ORFs ruled out from the comparative analysis because not in common between the tested datasets do not contain any of the reproducible profiles listed in Table S.4.

###### S.2.2.2 Analysing the reproducibility of Ribo-seq profiles for different genotypes

We reiterated our comparative analysis strategy to examine the role of the chosen *E. coli*'s genotype on the experimental reproducibility. Indeed, challenging the Samples of the groups D, E, F and G with our benchmark (group A Samples) we aimed at testing whether choosing the genotypes BW25113, BWDK or MC4100 instead of MG1655 might reduce the number of reproducible Ribo-seq profiles with respect to the 49 ones identified when the benchmark was analysed in isolation. Table S.8 summarises the results of this investigation. It turned out that the choice of the genotype is a variable to consider in terms of experimental reproducibility. Indeed, up to our results, while relying on BW25113 or BWDK genotypes do not impact significantly the experimental reproducibility with respect to the MG1655 genotype, when *E. coli* MC4100 (Samples G1$\alpha$, G2$\alpha$ and G3$\alpha$) is chosen, the number of reproducible Ribo-seq profiles falls to 28.

| **Group** | **Sample** | **Genotype** | **Tested ORFs** | **Reproducible Profiles** |
|---|---|---|---|---|
| D | D1$\alpha$ | E. coli k-12 BW25113 | 2238 | 47 |
| E | D1$\alpha$ | E. coli k-12 BW25113 | 2231 | 48 |
| F | F1$\alpha$ | E. coli k-12 BWDK | 2219 | 44 |
| G | G1$\alpha$ | E. coli k-12 MC4100 | 2226 | 28 |
| | G2$\alpha$ | E. coli k-12 MC4100 | 2226 | 28 |
| | G3$\alpha$ | E. coli k-12 MC4100 | 2226 | 28 |

Table S.8: Results of the comparative analysis to check the impact on experimental reproducibility of the chosen *E. coli*'s genotype. Column 1: Sample's group; Column 2: Sample's ID; Column 3: Sample's genotype; Column 4: number of tested ORFs, *i.e.*, number of ORFs in common between the tested Sample and the samples belonging to the benchmark group A; Column 5: number of reproducible profiles identified from the comparative analysis. **Note**: the ORFs ruled out from the comparative analysis involving group F because were not in common between groups A and F, included two of the 49 reproducible profiles identified from the analysis of the benchmark.

###### S.2.2.3 A set of resilient Ribo-seq profiles

From the intersection of the sets of genes that we characterised through the comparative analyses reported above, it is possible to identify a group of Ribo-seq profiles that result to be reproducible independently of either the colture medium or the *E. coli*'s genotype chosen for the analysis (Table S.9). Studying the biological properties of the corresponding genes or mRNAs would be surely interesting but it is beyond the scope of this paper. Nevertheless, it is worth to point out that these extremely resilient Ribo-seq profiles could represent a precious reference for normalisation when it might be needed for comparative purposes.

| Gene | Groups B1 B2 | Groups B3 B4 B5 | Groups C1 C2 | Groups C3 C4 | Group D1 | Group E1 | Group F1 | Group G1 |
|---|---|---|---|---|---|---|---|---|
| <u>*gltB*</u> | + | + | + | + | + | + | + | + |
| <u>*tufA*</u> | + | + | + | + | + | + | + | + |
| <u>*tufB*</u> | + | + | + | + | + | + | + | + |
| *hokB* | + | + | | | + | + | + | |
| *ubiJ* | + | + | + | + | | + | + | |
| <u>*nuoM*</u> | + | + | + | + | + | + | + | + |
| <u>*wbbH*</u> | + | + | + | + | + | + | + | + |
| <u>*wbbI*</u> | + | + | + | + | + | + | + | + |
| <u>*yfaD*</u> | + | + | + | + | + | + | + | + |
| <u>*rsxC*</u> | + | + | + | + | + | + | + | + |
| *gtrS* | + | + | | | + | + | | |
| rodZ | + | | + | + | + | + | + | |
| <u>ispB</u> | + | + | + | + | + | + | + | + |
| aceE | + | + | + | + | + | + | + | |
| alaS | + | + | | | | + | | + |
| ClpA | + | + | | | + | + | + | + |
| cyoA | + | + | | | + | + | + | |
| <u>DeaD</u> | + | + | + | + | + | + | + | + |
| dld | + | + | | | + | + | + | |
| dnaX | + | + | | + | + | + | + | |
| accC | + | + | + | | + | + | | + |
| gyrB | + | + | + | | + | + | + | |
| ileS | + | + | | | + | + | + | + |
| infB | + | + | | + | + | + | + | + |
| ligA | + | + | + | + | + | + | + | |
| lon | + | + | + | + | + | + | | |
| <u>MukB</u> | + | + | + | + | + | + | + | + |
| nagE | + | + | | | + | + | + | |
| <u>ompC</u> | + | + | + | + | + | + | + | + |
| parC | + | + | + | + | + | + | + | |
| pepN | + | + | | + | + | + | + | |
| secY | + | + | | | + | + | | + |
| pssA | + | + | | | + | + | + | + |
| rne | + | + | + | + | + | + | + | |
| rpoB | + | + | | | + | + | + | + |
| ftnA | + | + | | | + | + | + | |
| secA | + | + | | + | + | + | + | |
| sucA | + | + | | | + | + | + | + |
| thrA | + | + | | | + | + | + | + |
| <u>rnr</u> | + | + | + | + | + | + | + | + |
| waaO | + | + | | | + | + | + | |
| ubiA | + | + | | | + | + | + | |
| YifK | + | + | | | + | + | + | + |
| lptD | + | + | + | + | + | | + | + |
| proP | + | + | | | + | + | + | |
| <u>uup</u> | + | + | + | + | + | + | + | + |
| <u>ydbK</u> | + | + | + | + | + | + | + | + |
| dxs | + | + | | | + | + | + | |
| ycaO | + | + | | | + | + | + | + |

Table S.9: Set of 15 Ribo-seq profiles that resulted to be reproducible independently on *E. coli*'s genotype or the colture medium (Column 1, <u>underlined</u>). The entries in common with Table S.2 are *italicised*. Columns 1 and 2: benchmark set, i.e. genes of group A (Table S.5) associated with a high resolution Ribo-seq profile, after excluding the dataset A10$\zeta$. Columns 3 and 4: the genes belonging to the group $B$ and associated with reproducible Ribo-seq profiles after the comparison with the benchmark are marked with the symbol "+". Columns 5 and 6: the genes belonging to the group $C$ and associated with reproducible Ribo-seq profiles after the comparison with the benchmark are marked with the symbol '"+". Columns 7, 8, 9, 10: the genes belonging to the groups $D$, $E$, $F$ and $G$ respectively and associated with reproducible Ribo-seq profiles after the comparison with the benchmark are marked with the symbol "+".

### REFERENCES

#### References

[1] Barrett, T. *et al.* NCBI GEO: archive for functional genomics data sets—update. *Nucleic Acids Research* **41**, D991–D995 (2012).

[2] Edgar, R., Domrachev, M. & Lash, A. E. Gene Expression Omnibus: NCBI gene expression and hybridization array data repository. *Nucleic acids research* **30**, 207–10 (2002).

[3] Oh, E. *et al.* Selective ribosome profiling reveals the co-translational chaperone action of trigger factor in vivo. *Cell* **6**, 1295–1308 (2011).

[4] Woolstenhulme, C. J., Guydosh, N. R., Green, R. & Buskirk, A. R. High-Precision analysis of translational pausing by ribosome profiling in bacteria lacking EFP. *Cell Reports* **11**, 13–21 (2015).

[5] Morgan, G. J., Burkhardt, D. H., Kelly, J. W. & Powers, E. T. Translation efficiency is maintained at elevated temperature in E. coli. *The Journal of biological chemistry* jbc.RA117.000284 (2017).

[6] Mohammad, F., Woolstenhulme, C. J., Green, R. & Buskirk, A. R. Clarifying the Translational Pausing Landscape in Bacteria by Ribosome Profiling. *Cell Reports* **14**, 686–694 (2016).

[7] Li, G. W., Burkhardt, D., Gross, C. & Weissman, J. S. Quantifying absolute protein synthesis rates reveals principles underlying allocation of cellular resources. *Cell* **157**, 624–635 (2014).

[8] Subramaniam, A. R., Zid, B. M. & O'Shea, E. K. An integrated approach reveals regulatory controls on bacterial translation elongation. *Cell* **159**, 1200–1211 (2014).

[9] Guo, M. S. *et al.* MicL, a new $\sigma$E-dependent sRNA, combats envelope stress by repressing synthesis of Lpp, the major outer membrane lipoprotein. *Genes and Development* **28**, 1620–1634 (2014).

[10] Burkhardt, D. H. *et al.* Operon mRNAs are organized into ORF-centric structures that predict translation efficiency. *eLife* **6** (2017).

[11] Li, G.-W., Oh, E. & Weissman, J. S. The anti-Shine-Dalgarno sequence drives translational pausing and codon choice in bacteria. *Nature* **484**, 538–541 (2012).

[12] Kannan, K. *et al.* The general mode of translation inhibition by macrolide antibiotics. *Proceedings of the National Academy of Sciences* **111**, 15958–15963 (2014).

[13] Marks, J. *et al.* Context-specific inhibition of translation by ribosomal antibiotics targeting the peptidyl transferase center. *Proceedings of the National Academy of Sciences* **113**, 12150–12155 (2016).

[14] Hwang, J.-Y. & Buskirk, A. R. A ribosome profiling study of mRNA cleavage by the endonuclease RelE. *Nucleic Acids Research* **45**, 327–336 (2017).

[15] Haft, R. J. F. *et al.* Correcting direct effects of ethanol on translation and transcription machinery confers ethanol tolerance in bacteria. *Proceedings of the National Academy of Sciences of the United States of America* **111**, E2576–E2585 (2014).

[16] Baggett, N. E., Zhang, Y., Gross, C. A. & Ibba, M. Global analysis of translation termination in E. coli. *PLOS Genetics PLoS Genet* **13** (2017).
